# Molecular insights into the activation of Mre11-Rad50 endonuclease activity by Sae2/CtIP

**DOI:** 10.1101/2023.08.05.552090

**Authors:** Yoann Nicolas, Hélène Bret, Elda Cannavo, Petr Cejka, Valerie Borde, Raphael Guerois

## Abstract

In *S. cerevisiae*, the Mre11-Rad50-Xrs2 (MRX)-Sae2 nuclease activity is required for the resection of DNA breaks with secondary structures or protein blocks, while in humans, the MRE11-RAD50-NBS1 (MRN) homologue with CtIP is needed to initiate DNA end resection of all breaks. Phosphorylated Sae2/CtIP stimulates the endonuclease activity of MRX/N. Structural insights into the activation of the Mre11 nuclease are available only for organisms lacking Sae2/CtIP, so little is known how Sae2/CtIP activates the nuclease ensemble. Here, we uncover the mechanism of Mre11 activation by Sae2 using a combination of AlphaFold2 structural modeling, biochemical and genetic assays. We show that Sae2 stabilizes the Mre11 nuclease in a conformation poised to cleave substrate DNA. Several designs of compensatory mutations establish how Sae2 activates MRX *in vitro* and *in vivo,* validating the structural model. Last, our study uncovers how human CtIP, despite considerable sequence divergence, employs a similar mechanism to activate MRN.

## Introduction

DNA double-strand breaks (DSBs) represent a threat to cellular survival and genome stability. Cells possess two main DSB repair pathways: template-independent and potentially mutagenetic non-homologous end-joining (NHEJ) and template-directed, hence more accurate, homologous recombination (HR). HR is initiated by a process termed DNA end resection. In resection, the 5’- terminated DNA strand at a DSB site is nucleolytically degraded, resulting in the formation of a 3’- terminated single-stranded DNA (ssDNA) overhang. The overhang is then bound by the RAD51 strand exchange protein to form a nucleoprotein filament with the capacity to identify and pair with homologous DNA, which serves as a repair template. Once the DNA strand exchange takes place, the invaded 3’-terminated end can prime DNA synthesis. There are multiple HR sub-pathways leading to diverse genetic outcomes, however they all share the initial DNA end resection process ^1, 2^.

There are at least three conserved nucleases or nuclease complexes that catalyze DNA end resection in eukaryotic cells. Their usage and relative importance are dependent on the nature of the DNA break, cell cycle stage and physiological context. Resection pathways are generally divided into initial short-range and downstream long-range resection steps ^1^. Short-range resection is catalyzed by the Mre11/MRE11 nuclease, which functions within the Mre11-Rad50-Xrs2 (MRX) complex in the budding yeast model *Saccharomyces cerevisiae*, or MRE11-RAD50-NBS1 (MRN) in human cells ^3, 4^. Further downstream, either EXO1 or DNA2 nuclease extend the resection tracks to perform long-range resection ^3, 5, 6^. Beyond the initiation of DNA end resection, the MRX/N proteins also have nuclease-independent functions to promote NHEJ, or to activate DNA damage checkpoint through the control of the apical Tel1/ATM kinase ^7^.

MRN is generally required to initiate DNA end resection in human cells under all conditions, while MRX may be dispensable in budding yeast for the processing of chemically “clean” DSBs without protein adducts or secondary structures, such as hairpins ^8–10^. The unique feature of MRX/N is the ability to resect protein-blocked DSBs ^11^. Both yeast and human MRX/N help remove the NHEJ factor Ku from DNA ends, and in this way to channel non-productive NHEJ complexes into HR ^12–15^. MRX/N is also necessary to process meiotic DNA breaks, catalyzed in a programmed manner by Spo11 during the prophase of the first meiotic division ^16–18^. Spo11 remains covalently attached to the DNA end post-cleavage and needs to be removed by MRX before further processing and repair. The MRX/N complex employs a unique strategy, as it cuts the 5’-terminated DNA strand endonucleolytically some distance away from the DSB ^11, 18^. The incision site is thus located on the internal side of the protein block, and can serve as an entry site for the long-range resection nucleases acting downstream. The endonuclease activity of the MRX/N complex is strongly stimulated by a phosphorylated co-factor, Sae2 in yeast and CtIP in human cells ^11, 19^. Sae2/CtIP are phosphorylated in a cell cycle- and DNA damage-dependent manner ^20–23^. Particularly, cyclin-dependent kinase (CDK) phosphorylation of Sae2/CtIP at S267/T847 occurs in the S and G2 phases, which allows DNA end resection only in the cell cycle phases when sister chromatids are available as templates for repair. Mutations in components of the MRN complex or CtIP lead to genome instability and several human diseases ^24, 25^.

Due to the complex phenotypes associated with MRX/N defects, resulting from impaired recombination, end-joining and checkpoint signaling, biochemical and structural insights are necessary to uncover the underlying mechanisms of Mre11 nuclease activation. The Mre11-Rad50 proteins are heterodimers with a M_2_R_2_ stoichiometry ^26–29^. The nuclease, ATPase and DNA binding sites are a part of globular domains that form a base, from which protrude two Rad50 coiled-coils, joined by a Zn-hook at the apex of the structure ^29–31^. Biochemical assays revealed that in yeast phosphorylated Sae2 is bound by Rad50, albeit with a low affinity, and Xrs2 is largely dispensable for resection ^7, 23^. However, these studies did not explain how Sae2/CtIP activates the Mre11/MRE11 endonuclease. Attempts to obtain high resolution structures of the MRX/N complexes and Sae2/CtIP were unsuccessful to date, particularly as both MRX/N and Sae2/CtIP oligomerize, making the sample very heterogeneous for traditional structural biology approaches ^32^. Insights into a potential mode of Mre11 nuclease activation came from studies employing bacteria or archaea, which contain homologues of Mre11 and Rad50, but neither Sae2/CtIP nor Xrs2/NBS1 ^33–39^. From these studies, we learned that the MR homologues exist in at least two basic conformations. In the nuclease inactive, or resting state, ATP-bound Rad50 (SbcC) blocks the Mre11 (SbcD) active site, preventing the nuclease activity. Upon ATP hydrolysis, the complex undergoes a major structural change, which places Mre11 to the side of the dimer, enabling its endonuclease activity ^38^. Nevertheless, due to the lack of Sae2/CtIP in these organisms, it is not known how Sae2 participates in these structural transitions to activate the endonuclease of the MRX/N complex.

Here, we modeled the structures of Mre11-Rad50 in its various states, and provide insights into the mechanism of Mre11 endonuclease activation by phosphorylated Sae2. The data yielded structural details of key protein-protein interfaces that assemble the enzyme complex in the nuclease active state. The model was substantiated by the design of individual mutations that disrupt these interfaces, and of combinations of the mutations on each side of the three interfaces that compensate each other and allow the assembly of an active Mre11-Rad50-Sae2 ternary complex. These mutants were extensively characterized biochemically *in vitro* and in genetic assays that necessitate the action of the MRX nuclease, which validates the structural models. The data explain how Sae2 phosphorylation at the S267 CDK site activates the nuclease ensemble, and we also designed a Rad50 mutant that partially bypasses this activation. Sae2 and CtIP homologs are evolutionarily related although their sequence diverged drastically during evolution. The size of the homologs can vary from less than 200 amino acids in *P. tetraurelia* to more than 900 residues in *H. sapiens* ^40^, but also the sequence identity in the most conserved C-terminal region can be as low as 10% between *S. cerevisiae* and *H. sapiens* ^41^. Our analysis revealed how these protein interfaces evolved in higher eukaryotic systems and how, in human cells, CtIP is able to perform the same regulatory function despite its high sequence divergence with Sae2.

## Results

### Alphafold2 predicts two conformations for ScMre11 and ScRad50 complex consistent with an auto- inhibited and an activated state

Using AlphaFold2-multimer (AF2) algorithm, we predicted the structure of the (ScMre11)_2_-(ScRad50)_2_ assembly (M_2_R_2_) adapting the length of ScRad50 coiled-coil to match the size limits imposed by AF2. The models converged into two distinct classes of structures differing by a major conformational change at the interface between the Mre11 catalytic domains and the Rad50 dimer (Figure 1A and 1B). The class 1 model is similar to the auto-inhibited form of the nuclease as observed in the experimental structures of archaeal, bacterial and eukaryotic homologs (Figure 1A, Figure S1D) ^29, 34, 35^. In the auto- inhibited state, the Rad50 ATPase domain obstructs access to the Mre11 active site. Class 2 model places the catalytic domain of Mre11 in an entirely different orientation (Figure 1B, Figure S1E), which is highly asymmetric, and more similar to the arrangement experimentally observed with the active state of the *E. coli* EcMre11 homologs in the context of a DNA bound to the EcRad50 groove (Figure 1C) ^38^.

**Figure 1:**
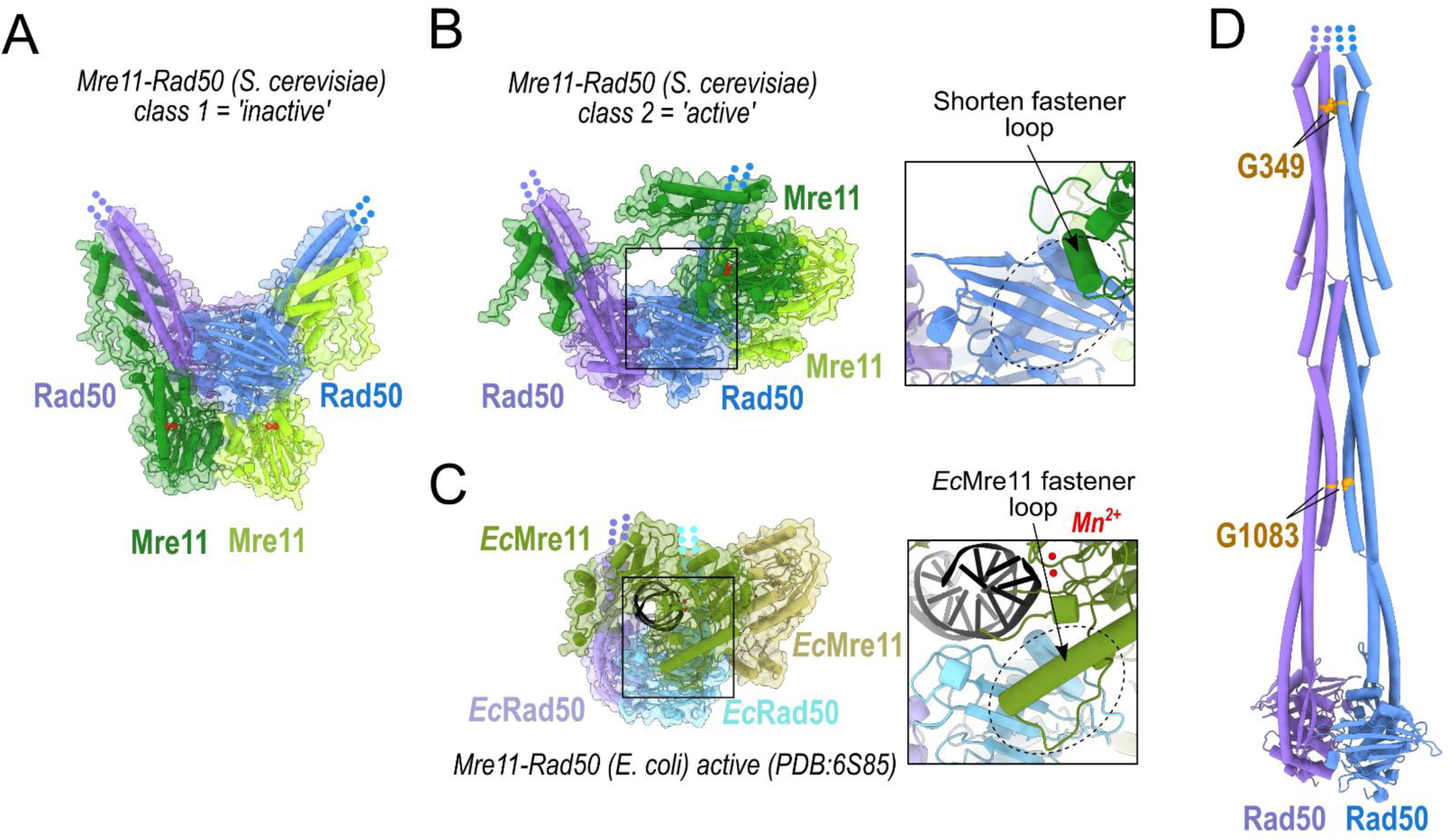
Structural modeling of *S. cerevisiae* Mre11-Rad50 assembly. (A) Structure of the *S. cerevisiae* Mre11 (1-533) (green) - Rad50 (1-215+1106-1312) (blue) complex generated in the first class of AF2 models in a 2:2 stoichiometry representative of an inactive form. (B) Structure of the same *S. cerevisiae* complex as in panel (A) generated in the second class of AF2 models more similar to an active form. The close-up view highlights that the fastener loop region is much shorter in eukaryotic Mre11 than in bacterial Mre11. (C) Experimental structure (PDB: 6S85) of the activated form of the *E. coli* Mre11-Rad50 complex in a 2:2 stoichiometry bound to a double-stranded DNA substrate. Close-up view of the fastener loop folding as a helix that stabilizes the active state. (D) Structural model of the *S. cerevisiae* Rad50 homodimer generated by AF2 (delimitations shown 1-400 and 950-1312), in which the coiled-coils assembled into a rod-like structure. The regions in contact between two coiled-coils involve highly conserved glycines (Figure S1A) that are labeled in brown at different positions along the coiled-coils. The G1083^ScRad50^ and G349^ScRad50^ are reminiscent of the G849^EcRad50^ in *E. coli,* whose mutation to alanine was shown to affect significantly the endonuclease activity of the bacterial enzyme ^38^.

In the *E. coli* active state MR complex bound to DNA, the coiled-coils of EcRad50 are wrapped around the DNA in a closed state, which brings the EcMre11 active site into the proper geometry to cleave dsDNA substrates. The closed conformation of the ScRad50 coiled-coils was only generated by AF2 for models integrating longer lengths of the Rad50 coiled-coil. Using a longer construct of (ScRad50)_2_ alone, embedding up to 711 residues out of 1312, and without adding ScMre11, we could generate a symmetrical structural model of ScRad50 coiled-coil forming a long rod using AF2 (Figure 1D, Figure S1A and S1C). This closed conformation with contact points regularly distributed along the coiled-coil sequence is similar to the rod structure observed in the low resolution cryoEM map of the *Escherichia coli* MR homolog ^38^ and in the recently reported structure of the *Chaetomium thermophilum* MRN complex ^29^ (Figure S1B). Thus, despite the different means of achieving MR nuclease activation between organisms, in particular with or without a Sae2/CtIP homolog, our multiple sequence alignment led AF2 models to converge on distinct classes of states likely to represent key evolutionarily encoded functional steps.

### Addition of ScSae2 to ScMre11 and ScRad50 favors the activated state conformation

In most eukaryotes, stimulation of the endonuclease activity of the MR complex requires the binding of a phosphorylated co-factor, Sae2 in *S. cerevisiae* or CtIP in *H. sapiens* ^11, 19^. To probe the mechanisms underlying this process, we predicted the structure of the complex formed between ScMre11, ScRad50 and ScSae2 using AF2. For simplicity, we omitted ScXrs2, because it is largely dispensable for DNA resection *in vitro* and *in vivo* ^7^. We also focused on the C-terminal domain of Sae2, which was previously shown to bind Rad50 ^23^. We did not include the N-terminal domain of Sae2 that is involved in tetramerization ^23, 42, 43^. Different constructs and lengths were explored ranging from a large (ScMre11)_2_-(ScRad50)_2_-ScSae2(180-345) (M_2_R_2_S) assembly to a smaller one as shown in Figure 2A (see Methods). In contrast to the (M_2_R_2_) assembly in which two classes of conformations were generated, the presence of ScSae2 generated only the class 2 (Figure 2A), resembling the activated form of the *E. coli* MR nuclease (Figure 1B and 1C), while the class 1, corresponding to the auto-inhibited form, was no longer generated. The stimulatory function of scSae2 may thus be explained by shifting the equilibrium of the MR structure towards the active conformation.

**Figure 2:**
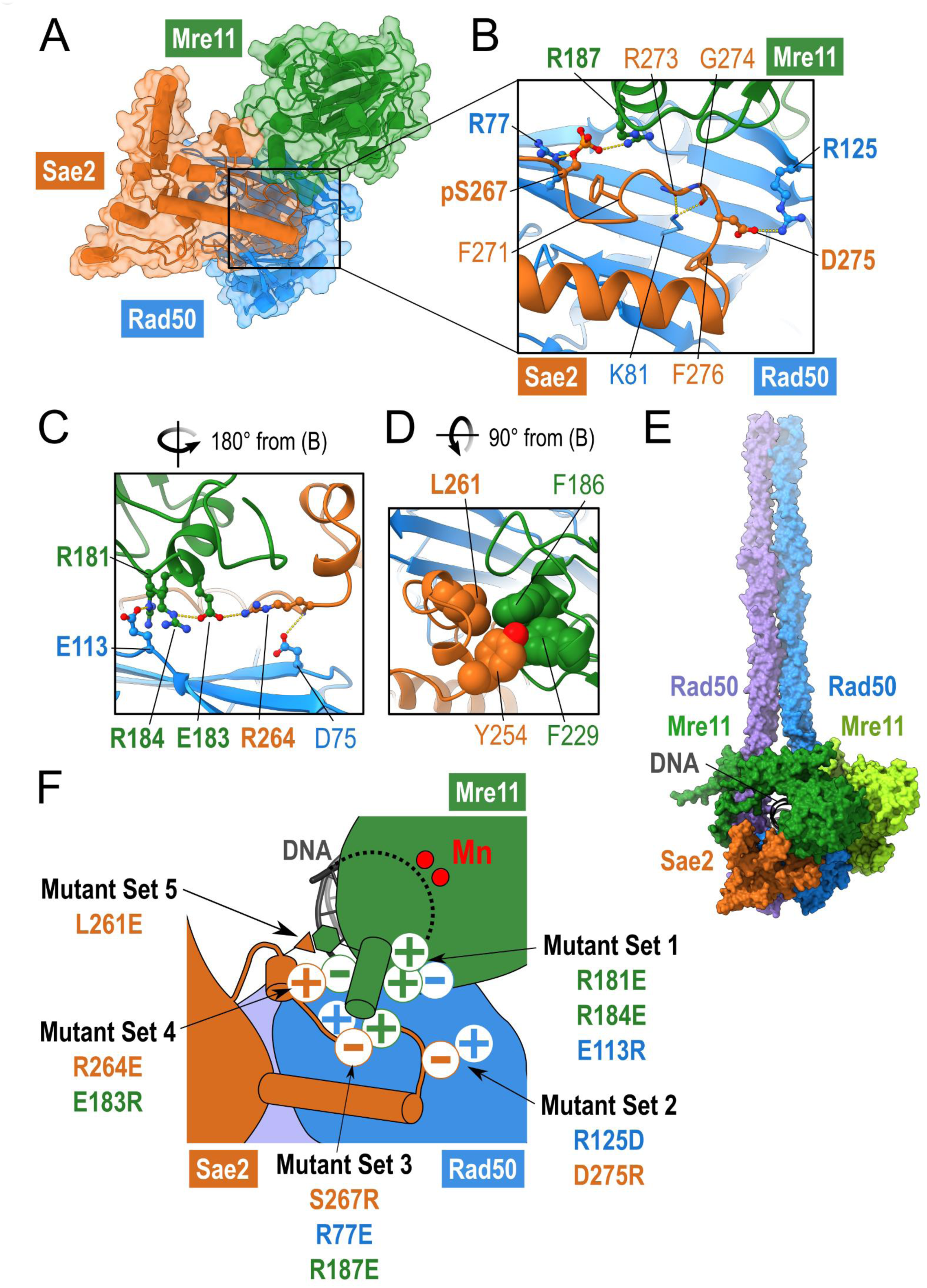
Structural modeling of *S. cerevisiae* Sae2-Mre11-Rad50 assembly. (A) Structural model of the *S. cerevisiae* Sae2 (180-345) (orange) - Mre11 (1-303) (green) - Rad50 (1- 166+1206-1312) (blue) complex generated by AF2 in a 1:1:1 stoichiometry representative of the active conformation form. (B) Close-up view of (A) highlighting the sidechains of interfacial residues forming an interaction between the loop of ScSae2 phosphorylated at S267. (C) Second close-up view corresponding to a different orientation with respect to panel (B) obtained after rotating the structure by 180° around a vertical axis and highlighting the network of salt-bridges formed at the interface of the trimer. (D) Third close-up view corresponding to a different orientation with respect to panel (B) obtained after rotating the structure by 90° around a horizontal axis and highlighting the cluster of apolar side-chains packing a the interface between ScSae2 and ScMre11. (E) General composite model generated by integrating models from Figure 1B, 1C, 1D and 2A of the *S. cerevisiae* Sae2 (180-345) (orange) - Mre11 (1-533) (green) - Rad50 (1-364+966-1312) (blue) in a 1:2:2 stoichiometry bound to DNA and shown as surface representation. (F) Summary of the five sets of disruptive and compensatory mutants designed to validate the importance of each of the contact points at the interface of the complex assembly.

### A ternary complex forms between ScSae2, ScMre11 and ScRad50

The structure of ScSae2 in its state bound to M_2_R_2_ (Figure 2A and Figure S2A) is distinct from any known fold in the PDB as established by the DALI server ^44^ (Table S1). In the model, the C-terminal domain of ScSae2 is predicted to bind one Rad50 monomer on its outer β sheet (Figure 2A). It also contacts the subunit of ScMre11 positioned to cleave dsDNA in the activated form, tightening the asymmetric arrangement of the Mre11 dimer on that of Rad50 (Figure 1C). Therefore, ScSae2 is predicted to create a trimeric interface connecting the outer β-sheet of ScRad50 and ScMre11 (Figure 2A and Figure S2D).

The region of ScMre11 in contact with ScSae2 and ScRad50 involves a short loop protruding from the catalytic domain and folded as a helix (Figure 2B). This ScMre11 helix makes only a few direct contacts with ScRad50 such as between residues E113^ScRad50^ and R181^ScMre11^/R184^ScMre11^ (Figure 2C, Figure S2B). In prokaryotes, a similar but larger helix, assigned as the fastener loop ^38^ (Figure 1C), was found in the EcMre11 homologue, and was shown to play an important role to connect EcMre11 and EcRad50 by binding the outer β sheet of Rad50 through a large network of polar and ionic interactions. Compared to the structure of the *E. coli* homologs, the interface of the eukaryotic ScMre11-ScRad50 subunits involving the helix exhibits a much smaller interaction surface (Figure 1B), suggesting that a part of the fastener function in eukaryotes was taken up by Sae2 (Figure S2B) ^39^. Indeed, the AF2 structural model of ScSae2 sheds light on this set of interactions, and Sae2 may strengthen the interaction of the small interface between ScMre11 and ScRad50 as will be seen below.

### Anchoring of ScSae2 to the ScRad50 outer β-sheet

Focus on the predicted interface between ScSae2 and ScRad50 identified an extensive network of interactions involving side-chain and backbone atoms of ScSae2 (Figure 2, Figure S2B). The most central residue involved in these interactions is S267^ScSae2^, which specifically needs to be phosphorylated for optimal stimulation of the MRX endonuclease ^11, 21^. The non phosphorylatable S267A^ScSae2^ mutant was shown to be inefficient in MR stimulation whereas phospho-mimicking S267E^ScSae2^ mutant partially retained the activity of phosphorylated wild-type Sae2. In our model, S267^ScSae2^ is positioned near R77^ScRad50^ enabling the formation of a salt bridge between both residues when S267^ScSae2^ is phosphorylated (Figure 2B). Downstream of S267^ScSae2^, another salt bridge between D275^ScSae2^ and R125^ScRad50^ was also predicted (Figure 2B). The K81^ScRad50^ ammonium moiety can form up to three hydrogen bonds with the carbonyl backbone atoms of the triad R273^ScSae2^-D275^ScSae2^. The involvement of K81^ScRad50^ is consistent with the original report of absence of meiotic DNA break processing, sporulation defects and spore lethality in cells bearing the K81I (termed *rad50S*) mutation^45^ (Figure 2B). This was later explained by the inability to remove Spo11 from break ends ^18^. In biochemical assays, the K81I mutation in ScRad50 was found to impair ScMre11 endonuclease activity by disrupting the physical interaction between phosphorylated ScSae2 C-terminus and ScRad50 ^11, 23^. Last, the ScSae2-ScRad50 interface is potentially further stabilized by the interaction of the highly conserved F271^ScSae2^ and F276^ScSae2^ aromatic rings packing against the carbon moieties of aforementioned R77^ScRad50^ and K81^ScRad50^ sidechains, respectively (Figure 2B).

### Inspection of the interface between ScSae2 and ScMre11 in the M_2_R_2_S ternary complex

Several contacts can be distinguished between ScSae2 and ScMre11, which likely favor the cooperative binding of ScSae2 and ScRad50 and the assembly with ScMre11 in a suitable geometry for nuclease activation. First, the phosphorylated form of S267^ScSae2^ can form a salt-bridge with R187^ScMre11^ (Figure 2B). Three residues upstream in ScSae2, a second salt-bridge can be formed between R264^ScSae2^ and E183^ScMre11^ located in the short helix protruding from ScMre11 (Figure 2C). The model also predicts the formation of a small hydrophobic cluster involving L261^ScSae2^ and F186^ScMre11^ with a potential contribution of surrounding side-chains from F229^ScMre11^ and Y254 ^ScSae2^ (Figure 2D), which likely further stabilizes the complex. Altogether, a larger network of interactions was found in the two interfaces involving ScSae2 with ScRad50 and ScMre11, while direct contacts between the Rad50 and Mre11 subunits appear rather limited compared to the complex formed in the bacterial homologs ^38, 39^. This property may explain the need for a central role of phosphorylated ScSae2 in stabilizing a functional and active nuclease. The global integration of the models shown in Figures 1B, 1D and 2A with the positioning of the DNA shown in Figure 1C could be achieved without any major structural rearrangement and converged on a coherent composite model (Figure 2E) representing the global architecture of the ScRad50-ScMre11-ScSae2 assembly and the DNA in the activated form of the complex.

### Assessment of the *S. cerevisiae* M_2_R_2_S structural model by targeted mutagenesis in biochemical and *in vivo* assays

Our structural model of the activated form of the ScM_2_R_2_-Sae2 predicts that phosphorylation of S267^ScSae2^ stabilizes a conformation of ScSae2 that promotes the positioning of ScMre11 on ScRad50 in its active state. To verify the predictions of this active structural arrangement, we designed 5 sets of mutants to probe the various interfaces predicted by AF2 (Figure 2F). We tested each mutation alone or in combination, to test for potential compensatory effects. To provide a simple quantitative readout for the compensatory effect of the mutations, we calculated the theoretical impact of the mutations assuming they were independent (i.e. on separate interfaces). Dividing the observed efficacy by the theoretical value gave us a rescue coefficient (R). A R value >1 suggests that the mutations compensate each other, and hence likely involve spatially close positions, while values ≈ 1 suggest no interaction.

We expressed and purified the wild-type and mutant recombinant proteins and protein complexes and tested for their capacity to endonucleolytically cleave protein-blocked DNA in reconstituted biochemical assays (Figure 3A and 3B and Figure S3). Additionally, we mutagenized *S. cerevisiae* cells and determined the effects of the individual and combined mutations *in vivo* in processes previously established to rely on the endonuclease activity of the MRX complex in conjunction with Sae2. To this point, we first employed the assay developed by Lobachev et al., which measures the recombination rate of DNA breaks terminated by hairpins formed by inverted repeated sequences, which require MRX-Sae2 activity for initial hairpin processing (Figure 3C) ^10^. Second, we probed for meiotic phenotypes. In *S. cerevisiae*, the Mre11 and Rad50 proteins are essential to promote DSB formation by the Spo11 transesterase, independently of the MRX nuclease activity and thus likely by a structural role (reviewed in ^17^). A complete loss of Mre11 or Rad50 therefore results in meiotic cells capable of sporulation, but forming only unviable spores, due to the absence of meiotic DSBs (see graph Figure 3D). However, the Mre11 nuclease activity is absolutely required for the removal of Spo11 oligonucleotides from meiotic DSB ends to allow for DSB resection and subsequent repair by HR, producing crossovers (Figure 3D). Mutants where the MRX is present but deficient in the Mre11 nuclease activity, such as *rad50^K81I^*(called *rad50S* ^45–47^) or *sae2*Δ, therefore allow meiotic DSB formation but exhibit greatly impaired sporulation efficiency and spore viability, reduced or absent Spo11 oligonucleotides and a strong reduction in meiotic crossovers.

**Figure 3:**
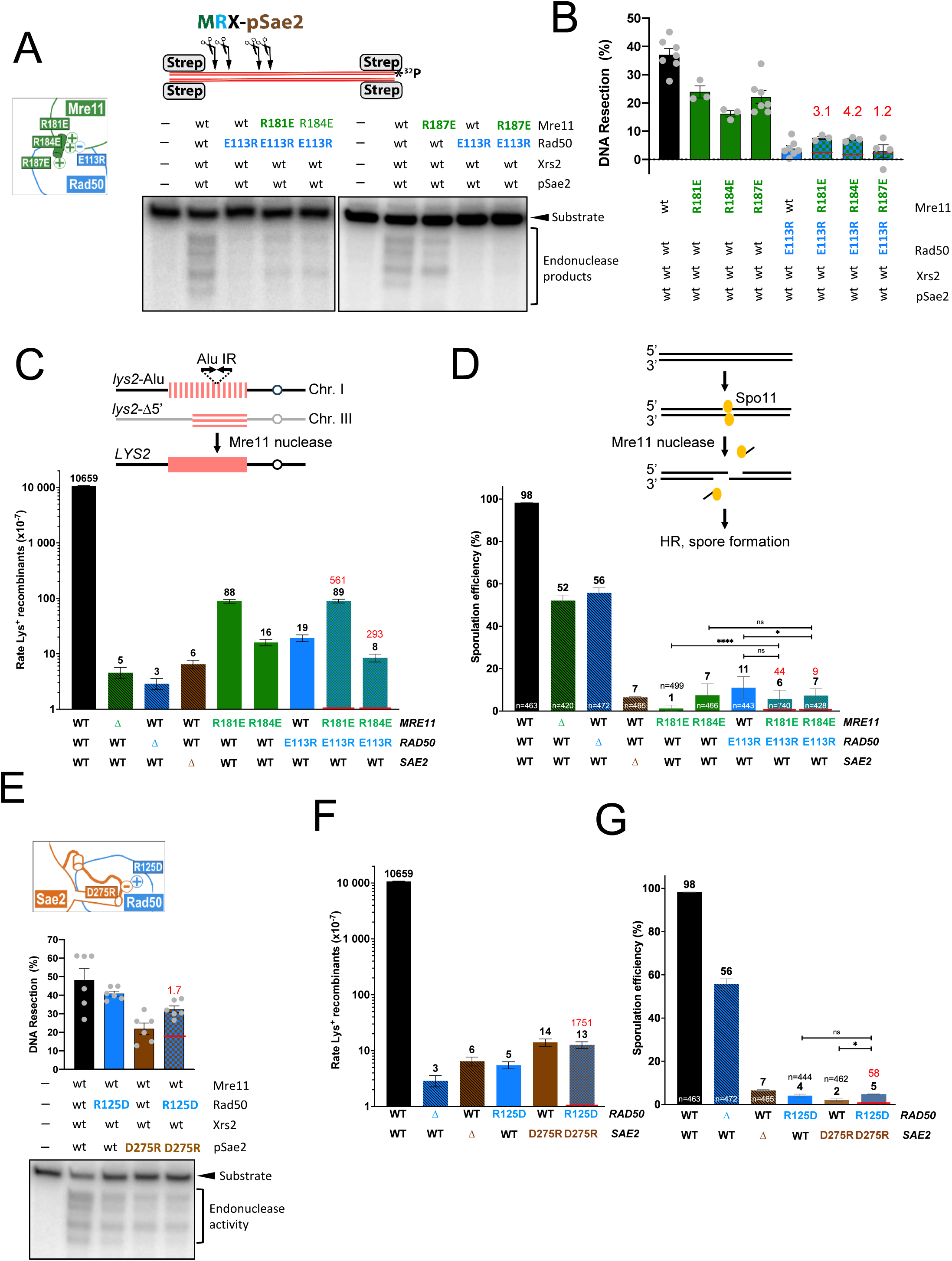
Validation of the *S. cerevisiae* M2R2S structural model by targeted mutagenesis. (A) Representative endonuclease assays with indicated proteins. Mre11 variants were co-expressed with Xrs2 (used at 25 nM), Rad50 variants were used at 50 nM, and pSae2 at 100 nM. (B) Quantitation of the assays such as shown in panel A. N=3-7; Error bars, SEM. The rescue coefficient (R), a value obtained by dividing the observed by the theoretical efficacy, assuming the mutations were independent, is shown in red. (C) Recombination frequencies of strains with the *lys2-AluIR* and *lys2-Δ5’* ectopic reporter system (illustrated by a cartoon). The rate of Lys+ recombinants is derived from the mean recombination frequency determined from six different isolates of each strain. Error bars represent confidence interval at 95% (see Methods). The red bar represents the expected recombination rate if we consider the two mutations as independent (i.e, if their effects were cumulative). The number in red indicates the rescue frequency (R), calculated as in (B). (D) Sporulation efficiency of diploid strains bearing the indicated Mre11 complex and *SAE2* homozygous alleles. As illustrated, a strain deficient for Mre11 nuclease does not process Spo11-deendent DSBs and shows high defect in sporulation. As in (A) and (C), the rescue coefficient R in the double mutants is indicated in red. Red bar: as in (C). Error bars represent the standard deviation (n=2). The total number of cells counted for each genotype is indicated. Statistical test: Fisher test. P-value ≥ 0,05 = NS ; P-value < 0,05 = * ; P-value < 0,01 = ** ; P-value < 0,001 = *** ; P-value < 0,0001 = ****. (E) Nuclease assays with the indicated variants (Mre11-Xrs2, 25 nM; Rad50, 50 nM; pSae2, 100 nM). Bottom, a representative experiment; top, a quantitation. N=6; Error bars, SEM.

Set 1 of mutants was designed to probe the predicted small interface between ScRad50 and ScMre11 as suggested by the model in Figure 2C. In this interface, R181^ScMre11^ and R184^ScMre11^ make a contact with E113^ScRad50^ (Figure 2C). We replaced the positively charged arginines R181^ScMre11^ and R184^ScMre11^ with negatively charged glutamic acid residues, creating R181E^ScMre11^ and R184E^ScMre11^ mutants. Reciprocally, on the Rad50 side, we generated the E113R^ScRad50^ charge reversal variant. *In vitro*, the E113R^ScRad50^ mutation alone strongly reduced the endonuclease activity (Figure 3A and 3B). The negative effects of the E113R^ScRad50^ mutation could be partially rescued by either R181E^ScMre11^ or R184E^ScMre11^, which had a moderate inhibitory effect in isolation (with ScRad50), with a rescue coefficient of 3.1 and 4.2, respectively. The observed rescue effect confirmed that the mutants impact the same interface, validating thus the structural model. In contrast, no rescue was observed in combination with R187E^ScMre11^, which is positioned at the other end of the scMre11 helix containing R181^ScMre11^ and R184^ScMre11^, and is thus less likely to interact with E113^ScRad50^ (rescue coefficient of 1.2), indicating no interaction.

The hairpin cleavage genetic assays exhibited a much higher dynamic range (Figure 3C). The single mutations E113R^ScRad50^, R181E^ScMre11^ and R184E^ScMre11^ reduced the recombination rates 560-, 120- and 660-fold with respect to the wild-type strain, respectively. The recombination rates of the double mutants between E113R^ScRad50^ and R181E^ScMre11^ or R184E^ScMre11^ were no worse than those of the single mutant strains, suggesting an interaction (rescue coefficients of 561 and 293, respectively). Finally, the meiotic phenotype, as assessed by the sporulation efficiency, confirmed these results, with the three single mutants strongly affected, to levels similar to the *sae2*Δ mutation, whereas the combination of E113R^ScRad50^ with R181E^ScMre11^ or R184E^ScMre11^ revealed rescue coefficients of 44 and 9, respectively, validating the relevance of the interaction also in the context of meiotic recombination (Figure 3D). Our biochemical and genetic experiments thus support the AF2 model predicting contacts between E113R^ScRad50^ and R181E^ScMre11^/R184E^ScMre11^.

Set 2 of mutants was designed to probe the ScRad50 and ScSae2 interface by evaluating the contribution of a potential salt bridge connecting R125^ScRad50^ and D275^ScSae2^ (Figure 2B). As above, we generated charge reversal mutations and evaluated the D275R^ScSae2^ and R125D^ScRad50^ mutants individually and combined. The rescue coefficient in biochemical experiments was rather modest (R=1.7) (Figure 3E), while in the genetic hairpin cleavage assays much more apparent (R=1751) (Figure 3F). Indeed, while the R125D^ScRad50^ mutation decreased recombination rate 760-fold in otherwise wild-type background, it increased recombination in the D275R^ScSae2^ cells, indicating a strong genetic interaction. Similarly, in meiosis, while both single mutants showed strongly decreased sporulation, similar to *sae2*Δ, the R125D^ScRad50^ mutation rescued the defect of D275R^ScSae2^ (R=58), confirming the strong genetic interaction (Figure 3G). These *in vivo* results support the structural AF2 model for the interface between ScRad50 and ScSae2.

### R77E substitution in Rad50 partially bypasses the requirement for co-activation of MRX by Sae2

Phosphorylation of the conserved S267 residue in Sae2 by cyclin-dependent kinase (CDK) strongly stimulates the endonuclease activity of MRX ^21,23^. While a non-phosphorylatable S267A^ScSae2^ substitution mutant is strongly impaired, the phosphomimetic S267E^ScSae2^ retains some activity and partially bypasses the requirement for CDK in resection ^21^. The AF2 model predicted that phosphorylated S267^ScSae2^ forms salt bridges with neighboring R187^ScMre11^ and R77^ScRad50^ (Figure 2B). In this way, phosphorylated S267^ScSae2^ promotes the positioning of the Mre11 and Rad50 subunits to specifically stabilize the nuclease in its active conformation. The third set of our mutants was designed to probe specifically this crucial trimeric interface. We designed S267R^ScSae2^; the positive charge of this normally negatively charged residue (in its phosphorylated form) was anticipated to confer a strong resection defect. We then set out to test this mutant in combination with charge reversal substitutions R187E^ScMre11^ and R77E^ScRad50^. As anticipated, the S267R^ScSae2^ mutant conferred a strongly impaired endonuclease activity to MRX *in vitro* (Figure 4A). A combination of S267R^ScSae2^ and R77E^ScRad50^ rescued resection *in vitro* (R=1.7) (Figure 4A). Likewise, the R187E^ScMre11^ and S267R^ScSae2^ mutations compensated each other *in vitro* (R=1.9). Hairpin cleavage assays verified these interactions in a cellular setting: the S267R^ScSae2^ substitution was as defective as the *sae2*Δ strain, and the R77E^ScRad50^ mutation increased recombination in S267R^ScSae2^ cells (R=1330) (Figure 4B). Likewise, a combination with R187E^ScMre11^ partially rescued recombination in S267R^ScSae2^ (R=552). Corroborating this effect on hairpin cleavage, in meiosis, the S267R^ScSae2^ mutant was as defective as the *sae2*Δ, and this defect was compensated by the R77E^ScRad50^ mutation, increasing sporulation efficiency (R=16) (Figure 4C), although not to a level sufficient to ensure spore viability (Figure S4). It is important to note that Spo11 catalyzes 150-200 DSBs in each meiotic *S. cerevisiae* cell. In order to achieve a full spore viability, all of these breaks have to be resected and repaired. Therefore, even a substantial rescue of DNA end resection, with only a few unrepaired breaks remaining, is not expected to result in spore viability. R187E^ScMre11^ also compensated the single S267R^ScSae2^, to a more modest extent, as measured by sporulation efficacy (R=9) (Figure 4C). These results converge towards S267^ScSae2^ closely and functionally interacting with R77^ScRad50^ and R187^ScMre11^.

**Figure 4:**
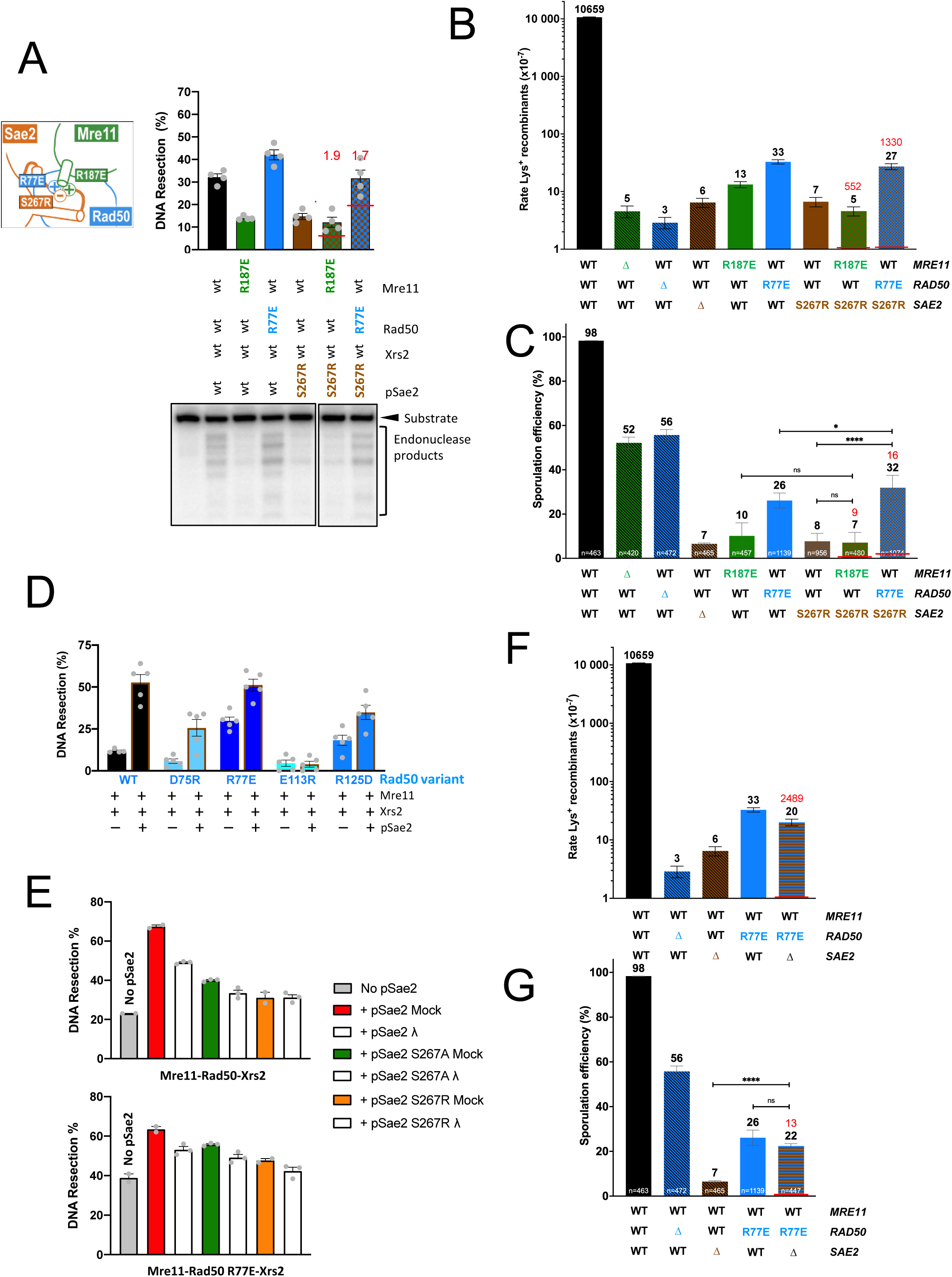
R77E substitution in Rad50 partially bypasses the requirement for co-activation of MRX by Sae2. (A) Nuclease assays with the indicated variants (Mre11-Xrs2, 25 nM; Rad50, 50 nM; pSae2, 100 nM). Bottom, a representative experiment; top, a quantitation. N=4; Error bars, SEM. (B) Recombination frequencies of strains with the *lys2-AluIR* and *lys2-Δ5’* ectopic reporter system, as in Figure 3C. (C) Sporulation efficiency, as in Figure 3D. (D) Quantitation of the nuclease assays with the indicated variants (Mre11-Xrs2, 25 nM; Rad50, 50 nM; pSae2, 100 nM). N=5; Error bars, SEM. (E) Quantitation of assays with either wild-type MRX (assembled from Mre11-Xrs2 heterodimer, 25 nM, and Rad50, 50 nM), or MR^R77E^X (assembled from Mre11-Xrs2 heterodimer, 25 nM, and Rad50^R77E^, 50 nM), and either mock-treated or λ phosphatase (λ) treated pSae2 variants. All pSae2 variants were prepared with phosphatase inhibitors, and hence were initially phosphorylated at multiple residues, before dephosphorylation by λ phosphatase, where indicated. N=3; Error bars, SEM. (F) Recombination frequencies of strains with the *lys2-AluIR* and *lys2-Δ5’* ectopic reporter system, as in Figure 3C. (G) Sporulation efficiency, as in Figure 3D.

Biochemical assays revealed that the R77E^ScRad50^ mutant was able to partially support resection without Sae2, more than any other variant constructed in this study (Figure 4D). Additionally, R77E^ScRad50^ promoted resection with pSae2 variants non-phosphorylatable at S267, but phosphorylated at other Sae2 sites, which is necessary to prevent protein aggregation (Figure 4E) ^23^. In *sae2*Δ cells, the recombination rate in the presence of R77E^ScRad50^ increased notably (R=2489), showing that R77E^ScRad50^ can partially bypass the requirement for Sae2 even in a cellular setting (Figure 4F). This was confirmed in meiosis, where R77E^ScRad50^ strongly compensated the *sae2*Δ mutant for sporulation (R=13) (Figure 4G). However, the functional rescue was likely not sufficient to ensure the repair of all meiotic DSBs, because spores remained unviable in the double mutant (Figure S4).

According to the AF2 model, the negative charge of R77E^ScRad50^ may place it in a favorable conformation with respect to positively charged R187^ScMre11^, and thus partially bypass the need for phosphorylated S267^ScSae2^ normally sandwiched in between. To our knowledge, R77E^ScRad50^ is the first reported mutation in the MRX complex that partially bypasses the requirement for Sae2 or its phosphorylation at the S267 site to co-activate its endonuclease activity *in vitro* and *in vivo*.

### Assessing the existence of a physical contact between ScSae2 and ScMre11

The AlphaFold2 model suggests that the ScSae2, ScMre11 and ScRad50 ternary assembly could be additionally cooperatively stabilized by direct contact between ScSae2 and ScMre11. Three residues upstream of S267^ScSae2^, we identified a salt bridge between the positively charged R264^ScSae2^ and negatively charged E183^ScMre11^ (Figure 2C) that could contribute to this tripartite stabilization. We set out to test a charge reversal mutation R264E^ScSae2^ in combination with E183R^ScMre11^. Both point mutants exhibited defects in the reconstituted endonuclease assay *in vitro*. A notable rescue effect was observed with combination of R264E^ScSae2^ and E183R^ScMre11^ *in vitro* (R= 1.8) (Figure 5A). In the hairpin cleavage recombination assay, the compensatory effect was quite dramatic (R= 2670, the strongest of any mutant combination in this study) (Figure 5B). Confirming these compelling *in vitro* and *in vivo* results, the meiotic sporulation efficiency defect of the E183R^ScMre11^ was almost entirely rescued by the R264E^ScSae2^ mutation, reaching nearly wild-type levels (R=10) (Figure 5C). This high rescue of sporulation prompted us to investigate if other readouts of meiotic recombination were also rescued. Indeed, this was the case, since spore viability of the double mutant was also nearly at a wild-type level, suggesting that most meiotic DSBs are repaired to ensure meiotic progression and viable spores (Figure 5D). Consistent with these dramatic compensatory effects, the amount of Spo11 oligonucleotides was nearly wild-type in the double mutant (Figure 5E). The combination of the biochemical and cellular assays validates the structural model that identified a close and specific interaction between the R264E^ScSae2^ and E183R^ScMre11^ residues, promoting the Mre11 nuclease.

**Figure 5:**
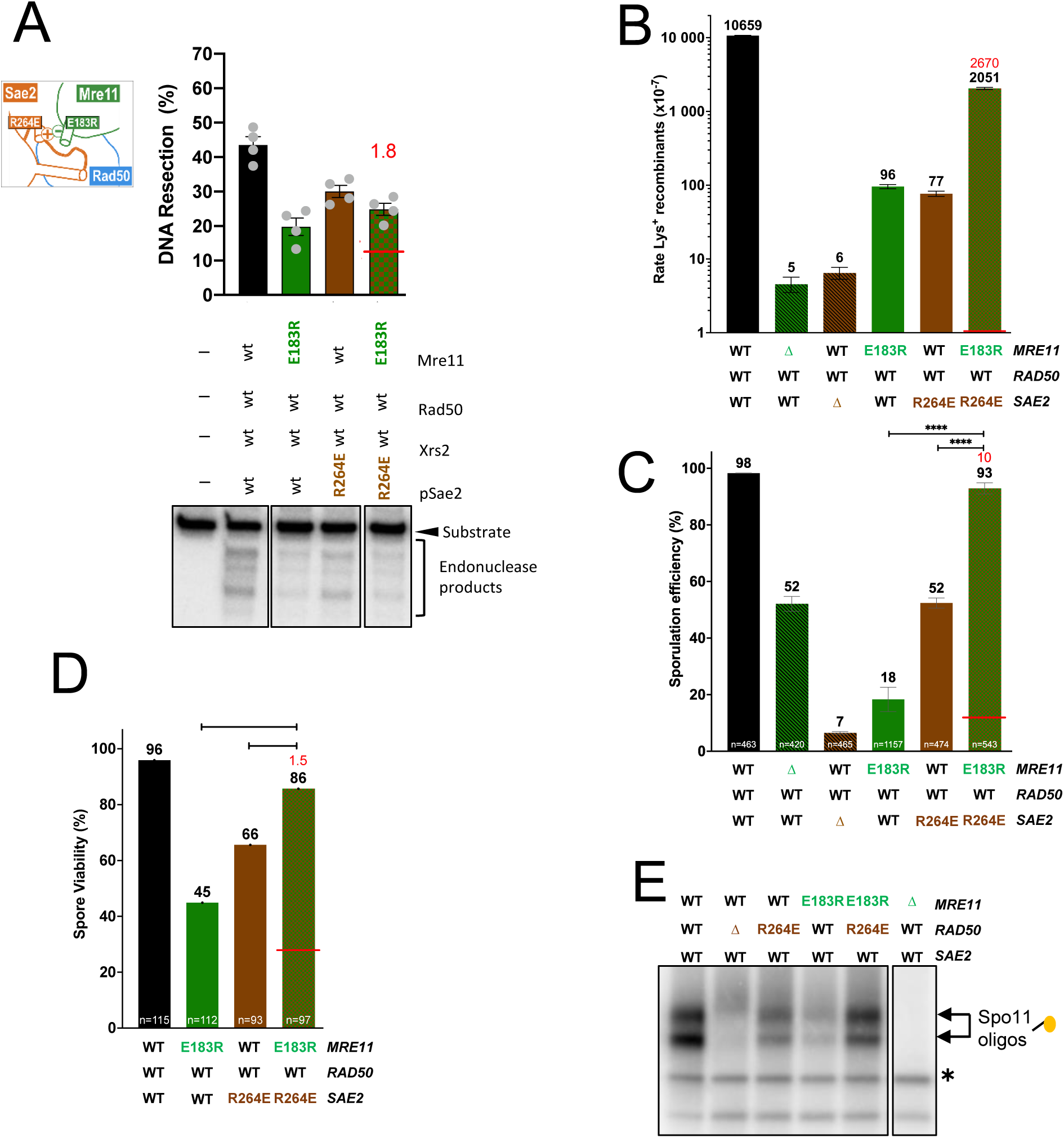
Validation of the interface between ScSae2 and ScMre11. (A) Nuclease assays with the indicated variants (Mre11-Xrs2, 25 nM; Rad50, 50 nM; pSae2, 100 nM). Bottom, a representative experiment; top, a quantitation. N=4; Error bars, SEM. (B) Recombination frequencies of strains with the *lys2-AluIR* and *lys2-Δ5’* ectopic reporter system, as in Figure 3C. (C) Sporulation efficiency, as in Figure 3D. (D) Spore viabilitity of strains with the indicated mutations. The number n of tetrad dissections is indicated for each. The statistical test used was a Fisher test between the observed and expected spore viability (P-value < 0,0001 = ****). Red bar and red number: as in Figure 3. (D) Spo11 oligonucleotide assay on strains with the indicated genotype. Two major bands correspond to Spo11 oligonucleotides. The asterisk marks a non-specific terminal deoxynucleotidyl transferase band.

### The small hydrophobic cluster between Sae2 and Mre11 favors the Mre11 nuclease

Finally, we tested the functional relevance of a hydrophobic cluster our model identified at the interface between Mre11 and Sae2, by mutating the L261^ScSae2^ residue, conserved in *Saccharomycetaceae* species (mutant 5, Figure 2F). In both the biochemical and *in vivo* meiotic assays, this mutant was defective, showing decreased *in vitro* resection (Figure S5A), as well a reduced sporulation (Figure S5B) and spore viability (Figure S5C). These results indicate that this hydrophobic patch supports optimal Mre11 conformation and nuclease activity.

## Discussion

In all eukaryotes, the nuclease activity of the MR machinery is strongly dependent on a conserved partner from the Sae2/CtIP family, whose sequences have diverged dramatically during evolution. The physical interaction between MR and Sae2 was reported to be weak and transient ^11,23^, likely due to the necessity to keep the MR endonuclease under a tight control. Nevertheless, the AlphaFold2 generated models reported here predict a ternary complex formation that was validated by mutagenesis in a combination of biochemical experiments *in vitro* and genetic assays in both vegetative and meiotic yeast cells.

Our structural, biochemical and genetics study in *S. cerevisiae* revealed that the transition from the inactive state of Mre11 to its active state, is triggered by the establishment of a cooperative set of interactions centered on the phosphorylated status of S267^ScSae2^ (Figure 6). The addition of a negative charge on S267^ScSae2^ by CDK-mediated phosphorylation switches the balance between unfavorable and favorable interactions that can form at the outer β-sheet of Rad50 with Mre11. In ScSae2, the conserved residues L261^ScSae2^, R264^ScSae2^ and D275^ScSae2^ provide key complementary contributions to the stabilization of the tripartite complex with ScMre11 and ScRad50. The validated structural model provides further insights on how Mre11 could be released from its auto-inhibitory conformation (Figure 1A) upon S267^ScSae2^ phosphorylation. In the Sae2 bound state, the backbones of residues R320- Q322 (in helix α9) (Figure S6A) are predicted to induce a steric clash with the capping domain of ScMre11 in its inactive state, thus favoring its relocation toward the active state. Interestingly, the telomeric protein Rif2 was proposed to antagonize this active conformation to specifically inhibit the endonuclease activity of MRX at telomeres ^48–50^.

**Figure 6:**
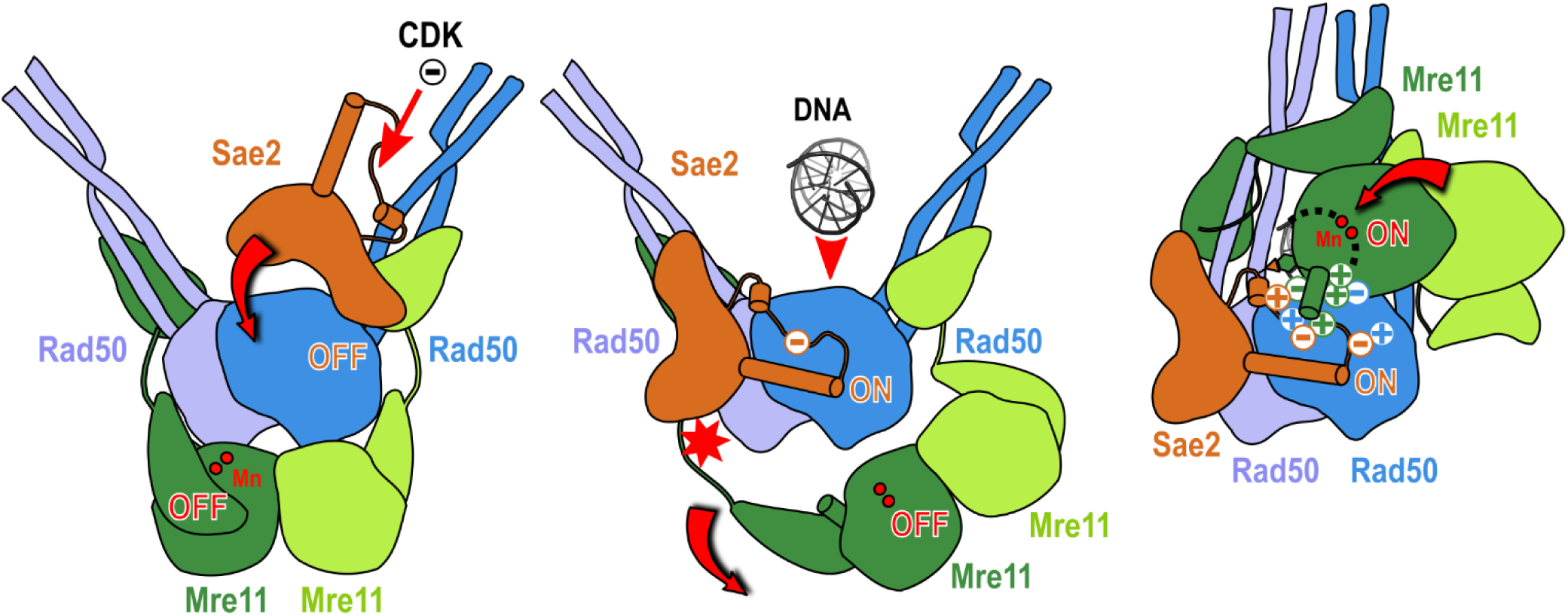
Schematic model for the molecular mechanism of activation of the nuclease complex. General model of the activation of the *S. cerevisiae* Mre11-Rad50-Sae2 complex upon phosphorylation of S267 by CDK. In step 1, the states of Sae2 and Mre11 are off. In step 2, binding of the phosphorylated loop to the outer β-sheet is incompatible with Mre11 in off-state. The added negative charge is represented by a circle with a minus sign and the red star reports steric hindrance. In step 3, one of the Mre11 dimer forms a trimeric assembly positioning Mre11 catalytic nuclease site in a position poised to cut DNA (represented by Mn atoms as yellow circles).

Sae2 and CtIP homologs are evolutionarily related although their sequence diverged drastically during evolution. Intriguingly, the residues L261^ScSae2^, R264^ScSae2^ and D275^ScSae2^ whose importance for the formation of the tripartite complex with Mre11 and Rad50 have been shown, are essentially conserved in the small subset of *Saccharomycetaceae* species (Figure 7A and Figures S2C and S7D) questioning the generality of the unraveled mechanisms. In ScSae2, the C-terminal domain adopts a combination of two small β-strands packing against α2 and α7-α9 helices (Figure S2A) that seems completely absent in human CtIP (Figure 7A). On another hand, in most eukaryotes CtIP contains a highly conserved “CxxC…RHR” motif (termed C-rich-H domain) that is completely absent in ScSae2 and in related species (Figure 7A). Interestingly, in *Yarrowia lipolytica*, a yeast belonging to the same subphylum as *S. cerevisiae* (*Ascomycotina*), the features specific to either ScSae2 (2 β-strands domain) or HsCtIP (C- rich-H domain) appear to coexist in the Sae2/CtIP hybrid ortholog, noted YlCtIP (Figure 7A).

**Figure 7:**
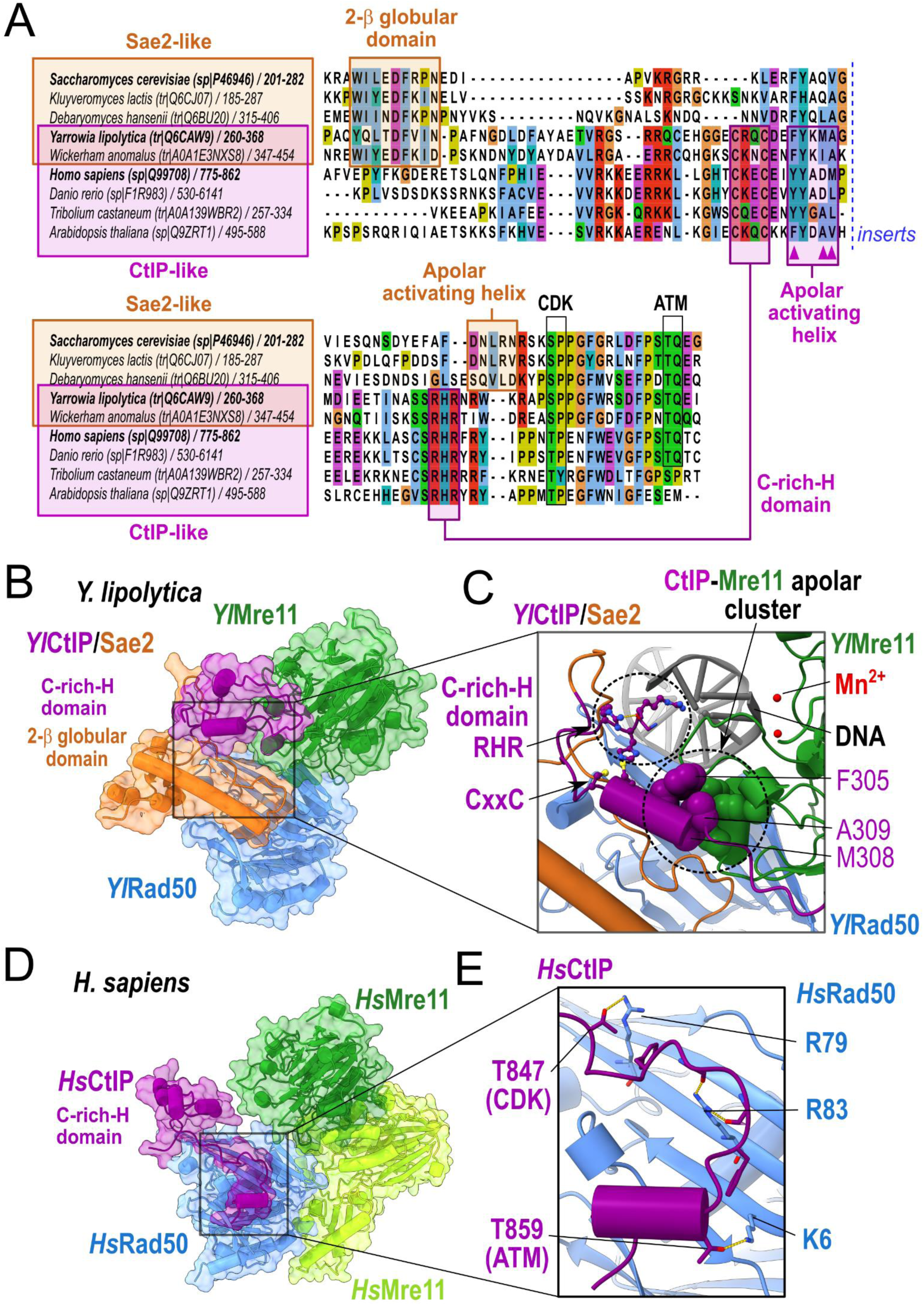
Evolution at the sequence and structural level of the interactions between Sae2/CtIP orthologs and the Mre11-Rad50 complex. (A) Multiple sequence alignment of Sae2 and CtIP orthologs in the conserved C-terminal domain involved in the Mre11 nuclease activation. Orange boxes indicate species with a 2 β-stranded globular domain and the conserved positions associated with the presence of this domain. Purple boxes indicate species with a cysteine-rich and histidine domain in their Sae2/CtIP orthologs and the positions defining this domain. Two boxes indicate the positions of the helical segment that the models identify as contributing to the formation of the apolar cluster between Sae2 or CtIP and Mre11 in their active conformations. Species in bold correspond to those for which a structural model of the complex with Mre11 and Rad50 was generated and analyzed. Purple triangles indicate the positions involved in the formation of apolar clusters with Mre11 in the active conformation in the orthologs having a C- rich-H domain. (B) Structural model of the *Y. lipolytica* CtIP/Sae2 (255-407) (orange+purple) - Mre11 (1-296) (green) - Rad50 (1-167+1179-1292) (blue) complex generated by AF2 in a 1:1:1 stoichiometry representative of the active conformation form. (C) The close-up view of the model from panel B is rotated by 90° with respect to panel B and highlights the relative orientation of the RHR, CxxC and apolar cluster motifs when the C-rich-H domain interacts with the Mre11 fastener helix in the active form. (D) Structural model of the *H. sapiens* CtIP (792-869) (purple) – 2 Mre11 (1-304 and 1-515) (green) - Rad50 (1-220+1098-1312) (blue) complex generated by AF2 in a 1:1:1:1 stoichiometry representative of the active conformation form. (E) Close-up view of the model from panel D showing the sidechains of the threonine residues of CtIP phosphorylated by CDK and ATM and which interact with basic sidechains of Rad50 outer β–sheet surface.

A model of the structure of the *Y. lipolytica* Rad50-Mre11-CtIP complex generated with AF2 provides key insights to understand how the C-rich-H domain could contribute to the activation of the MR nuclease. The C-rich-H domain is predicted to fold into a small two-helix domain, whose geometry is constrained by the highly conserved cysteine and histidine residues forming a cluster (Figure 7B and Figures S7A and S7B). The structural arrangement of the YlMR-CtIP ternary complex is very similar to that obtained for the ScMR-Sae2 complex. Of utmost interest, as in the structure of ScMR-Sae2, an apolar cluster is also formed between YlCtIP and YlMre11 involving the short fastener helix of YlMre11 (Figure 7C). In the case of *Y. lipolytica*, one of the helices of the C-rich-H domain mediates this interaction by exposing conserved apolar residues that pack against aromatic residues of the YlMre11 fastener helix, as in the MR-Sae2 complex (purple triangles in Figure 7A). In *S. cerevisiae*, deprived of C-rich-H domain, this interaction was in part mediated by L261^scSae2^ as measured by ‘mutant set 5’ assays (Figure 2D and Figure S5). Thus, the model of the *Y. lipolytica* assembly suggests that a role of the highly conserved C-rich-H domain is to create an anchor between Mre11 and CtIP to stimulate the active conformation of the nuclease. This orientation would additionally place conserved basic residues in favorable position to interact with DNA.

We further compared the evolutionary properties of Sae2 and CtIP orthologs by performing AF2 structural predictions between human HsRad50, HsMre11 and HsCtIP (Figure 7D and Figure S7C). AF2 models of the human assembly lead to a conformation between the C-rich-H^HsCtIP^ domain and HsMre11 that was not as compact as for the *Y. lipolytica* proteins, but the two domains come close to each other in a similar orientation. Conserved hydrophobic residues also protrude from the C-rich-H^HsCtIP^ and could also favorably make contacts with the aromatic residues of the fastener HsMre11 with little conformational change required. The contribution of DNA or other unmodeled partners in the assembly may be necessary to reach the fully active geometry with the human proteins. Just downstream of the RHR motif, T847^HsCtIP^, equivalent to S267^ScSae2^, is also phosphorylated by CDK ^22^. In this phosphorylated loop, the architecture of the state bound to the outer β sheet of HsRad50 is very similar to that modeled for the *S. cerevisiae* complex, with T847^HsCtIP^ contacting R79^HsRad50^, an arginine in the position equivalent to R77^ScRad50^ (Figure 7E; compare with Figure 2B). The possibility of establishing a charged salt-bridge interaction upon phosphorylation may therefore explain the stabilization of the MR-CtIP complex as in ScSae2. In humans, phosphorylation on another residue, T859^HsCtIP^, by ATM/ATR kinase was shown to also promote MR activity ^51^. In the structural model, T859^HsCtIP^ contacts a basic residue, K6^HsRad50^, which may additionally support the interaction with Rad50 upon CtIP phosphorylation (Figure 7E). The variety of ways in which the physico-chemical complementarity is encoded at the interface between the homologs of Sae2/CtIP, Rad50 and Mre11 is also highlighted when looking at the case of SpCtp1, the ortholog of HsCtIP *in Schizosaccharomyces pombe*. A specific polypeptide containing the conserved 15 amino acids at the C terminus of SpCtp1 is sufficient to stimulate Mre11 endonuclease activity in the absence of phosphorylation by CDK ^52^. In SpCtp1, the position corresponding to the phosphorylated residues S267^ScSae2^ and T847^HsCtIP^ is occupied by a hydrophobic amino-acid I283^SpCtp1^. On the other hand, in SpRad50, the position equivalent to the basic residues R77^ScRad50^ and R79^HsRad50^ is a L77^SpRad50^. This physico-chemical switch from an electrostatic to an apolar interaction could account for Ctp1 stabilizing the interaction with SpRad50 in a CDK-independent manner and promoting the formation of a tri-partite interface with SpMre11 to activate its endonuclease activity.

In conclusion, the comparative analysis of several interologous complexes has provided key insights into functional configurations and mechanisms for interpreting the role of the conserved motifs such as the C-rich-H domain in CtIP homologs. The CtIP and Sae2 families appear to have undergone considerable structural reorganization during evolution, while maintaining an ability to stabilize the active conformation of the Mre11 nuclease domain so that it can act on DNA clamped by Rad50.

## Acknowledgments

We thank Kirill Lobachev (Georgia Tech University) for the hairpin assays yeast strains, and Matt Neale (Sussex University) for advice on the Spo11 oligonucleotides protocol. This work was funded by Agence Nationale de la Recherche (ANR-21-CE44-0009) to R. G. and V. B. and Fondation pour la Recherche Médicale (Equipe FRM EQU201903007785) to V.B. The Swiss National Science Foundation (SNSF) (Grants 310030_207588 and 310030_205199) and the European Research Council (ERC) (Grant 101018257) support the research in the Cejka laboratory.

## Author Contributions

R.G., P.C. and V.B. designed the study. H.B. and R.G. performed the structural modeling and designed all the mutations. E.C. and P.C. performed all biochemical experiments. J.N. and V.B. designed and carried out the genetic assays in yeast. R.G., P.C. and V.B. wrote the manuscript. All authors contributed to prepare the final version of the manuscript.

## Methods

### Structural modeling

Sequences of *S. cerevisiae*, *Y. lipolytica* and *H. sapiens* for the Mre11, Rad50 and Sae2/CtIP were retrieved from UniProt database ^53^ (Table S2). Mre11, Rad50 and CtIP from *H. sapiens* or *Y. lipolytica* were used as input of mmseqs2 homology search program ^54^ to generate a multiple sequence alignment (MSA) against the UniRef30 clustered database ^55^. Homologs sharing less than 25% sequence identity with their respective query and less than 50% of coverage of the aligned region were not kept. For Sae2, we used HHblits ^56^ for its enhanced sensitivity with 8 iterations to retrieve remote homologs against the UniRef30_2020_06 database. In every MSA, in case several homologs belonged to the same species, only the one sharing highest sequence identity to the query was kept. Full-length sequences of the orthologs were retrieved and re-aligned with mafft ^57^. To model the structures of the Mre11-Rad50-Sae2/CtIP complexes, the corresponding MSAs were concatenated. In the concatenated MSAs, when homologs of different subunits belonged to the same species, they were aligned in a paired manner, otherwise they were left unpaired ^58^. The number of paired and unpaired sequences in the final concatenated MSAs are reported in Table S2. Concatenated MSAs were used as inputs to generate 25 structural models (5 different seeds) for each of the conditions using a local version of the ColabFold v1.3 interface ^59^ running 6 iterations of the Alphafold2 v2.2.0 algorithm ^60^ trained on the multimer dataset ^61^ on a local HPC equipped with NVIDIA Ampere A100 80Go GPU cards. Four scores were provided by AlphaFold2 to rate the quality of the models, the pLDDT, the pTMscore, the ipTMscore and the model confidence score (weighted combination of pTM and ipTM scores with a 20:80 ratio) ^61^. The scores obtained for all the generated models are reported in Table S3 for the model ranked first according to the confidence score. The models ranked first for each complex were relaxed using rosetta relax protocols to remove steric clashes ^62^ under constraints (std dev. of 2 Å for the interatomic distances) and were deposited in the ModelArchive database (https://modelarchive.org/) (Table S4). The global composite model of the (Mre11)2-(Rad50)2-Sae2 in complex with DNA was built by combining the *S. cerevisiae* models of (i) the long Rad50 dimer (Figure 1D, ModelArchive ID: ma- i9sv6), (ii) the Mre11 dimers in the active conformation with Rad50 (Figure 1B, ModelArchive ID: ma- jfoh4) and (iii) the trimeric assembly between Sae2, Mre11 and Rad50 (Figure 2A, ModelArchive ID: ma-qhmpy). Model (iii) was first superimposed on model (i) to recover the relative position of Sae2 and of one Mre11 nuclease domain in the context of the closed Rad50 coiled-coil, taking the outer β- sheet of Rad50 binding to Sae2 as a reference. The Mre11 nuclease domain from (iii) was then taken as reference to model the relative position of the second Mre11 nuclease domain using model (ii). The orientation of Mre11 (1-412) regions were then kept fixed while Mre11 domains (429-533) binding to Rad50 coiled-coils as in model (ii) were modeled on the closed conformation of the Rad50 coiled-coil using model (i) as reference for superposition. The flexible linkers Mre11 (412-429) connecting the two Mre11 domains of each subunit were finally remodeled using RCD+ ^63^. The positions of DNA, nucleotides and Mn^2+^ ions were modeled using the superposition of the composite global model with the bacterial assembly in PDB 6S85 taking the position of the Rad50 ATPase domains as a reference. The resulting global model was relaxed using rosetta under the same conditions as described above. The Molecular graphics and analyses were performed with UCSF ChimeraX ^64^. Multiple sequence alignments were represented using JalView ^65^.

### Yeast strains

*S. cerevisiae* strains used for all the meiotic experiments are diploids of the efficiently sporulating SK1 background ^66^. Strains used for the hairpin-opening assays are derived from the haploid HS21 strain, of the CGL background ^10, 67^. Their genotypes are listed in Table S5. Cells were grown and all assays were performed at 30°C. Strains were transformed by the lithium acetate method. Spo11-His6-Flag3::NatMX allele was amplified by PCR. Haploid *mre11*Δ, *rad50*Δ and *sae2*Δ strains for hairpin cleavage assays were provided by K. Lobachev. *mre11*Δ and *sae2*Δ in the SK1 diploid strains were obtained by replacing the entire coding sequence by the KanMX resistance marker ^68^. Introduction of all the *mre11*, *rad50* or *sae*2 point mutations were done in two steps: first, the regions surrounding the aminoacids to mutate were replaced by the KanMX cassette by PCR with long flanking homology. The regions deleted were *mre11Δ113-291*; *rad50Δ1-192* (which we also used as the “*rad50Δ*” for the meiotic experiments, since it deletes the start codon and the first 192 residues of the protein); *sae2Δ193-345*. Then, these strains were co-transformed with a NatMX selectable plasmid encoding both Cas9 and a guide RNA against the KanMX sequence and a mutagenized PCR fragment spanning the deletion as a healing fragment. Transformants were selected on clonNAT, and all transformants were sequenced for correct introduction of the mutation and preservation of the remaining of the targeted gene.

### Yeast hairpin cleavage assays

The rate of Lys+ recombinants was computed as the number of colonies growing on -Lys plates compared to the total number of colonies growing on YPD rich medium ^10^. The mean recombination rate was calculated from six different colonies of each strain using the Falcor website tool using the MSS-MLE ^69^ and (https://lianglab.brocku.ca/FALCOR/index.html).

### Yeast meiotic assays

Meiosis was induced synchronously by pre-growing cells in SPS pre-sporulation medium as previously described ^70^. Sporulation efficiency was measured after 24 h in sporulation medium as the percentage of cells that showed cells with two or more spores. At least two independent experiments were performed and at least 400 cells were counted for each genotype. For spore viability, cells were put on solid SPO plates for 48 h at 30°C. Spo11 oligonucleotide analysis was performed by immunoprecipitating Spo11-His6-Flag3 cells at t = 4hr in sporulation medium essentially as described in ^71^, with the following modifications: Dynabeads Protein G beads (Thermo Fisher Scientific : 10003D) and mouse anti-FLAG M2 monoclonal antibody (Sigma-Aldrich : F3165) were used. Samples were run on an Invitrogen NuPAGE Novex Bis Tris gel 4-12% gradient SDS-PAGE gel and dried under vacuum before exposing to a Phosphorimager screen.

### MRX and Sae2 purification

All proteins were expressed in *Spodoptera frugiperda* 9 (*Sf*9) cells. The Mre11 construct (vector pTP391 provided by T. Paull) contained a His-tag at the C- terminus, and was co-expressed and co-purified with Xrs2, tagged at the C-terminus with FLAG (pTP694 vector from T. Paull). The heterodimer was purified using affinity chromatography using NiNTA resin (Qiagen) followed by anti-FLAG M2 affinity resin (Sigma) ^11^. Rad50, FLAG-tagged at the C-terminus (vector pFB-Rad50-FLAG), was purified by using anti-FLAG M2 affinity resin (Sigma) ^72^. Phosphorylated Sae2 (pSae2), MBP-tagged at the N-terminus and his-tagged at the C-terminus, was prepared as described previously in the presence of phosphatase inhibitors ^23^ (vector pFB-MBP-Sae2-his). The MBP tag was cleaved and removed during protein purification. The λ phosphatase treatment was carried out as described previously ^23^. Point mutagenesis of the expression vectors was carried out using the QuikChange site directed mutagenesis kit (Agilent) according to standard procedures. The mutant proteins were then expressed and purified in the same way as the wild type proteins.

### Endonuclease assays

Nuclease assays (15 μl volume) were carried out with oligonucleotide-based DNA substrate blocked at all ends with streptavidin (1 nM molecules, oligonucleotides, 210: (GTAAGTGCCGCGGTGCGGGTGCCAGGGCGTGCCCTTGGGCTCCCCGGGCGCGTACTCCACCTCATGCATC and 211: GATGCATGAGGTGGAGTACGCGCCCGGGGAGCCCAAGGGCACGCCCTGGCACCCGCACCGCGGCACTTAC)^11^, in a buffer containing 25 mM Tris-acetate, pH 7.5, 1 mM dithiothreitol, 5 mM magnesium acetate, 1 mM manganese acetate, 1 mM ATP, 80 U/ml pyruvate kinase (Sigma), 1 mM phosphoenolpyruvate, 0.25 mg/ml bovine serum albumin (New England Biolabs). 30 nM streptavidin (Sigma), was added to the substrate containing the reaction mixture and incubated for 5 min at room temperature. Recombinant proteins were then added on ice and incubated for 30 min at 30°C. Reactions were terminated and the DNA products were separated by denaturing gel electrophoresis and detected by autoradiography ^73^. The data were quantitated using Fiji (Image J). The signal below the substrate band in no protein lane was removed from all lanes as a background. DNA end resection was then calculated as the fraction of the endonucleolytic DNA cleavage products in each lane. The data were plotted using GraphPad Prism (V8).

## Supplementary material

### Supplementary Figures legends

**Figure S1:**
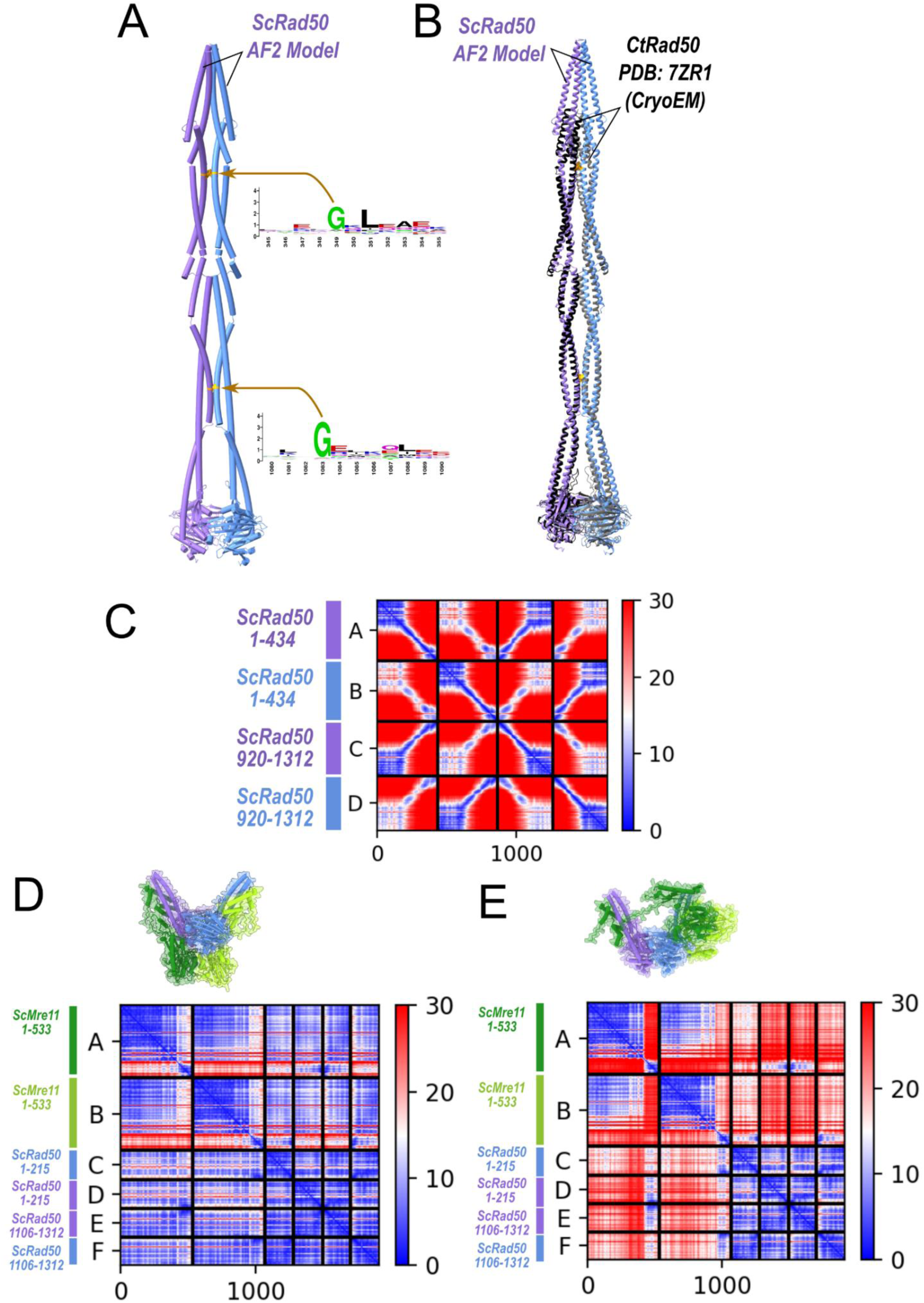
Details on the structural modeling of *S. cerevisiae* Mre11-Rad50 assembly. Related to Figure 1. (A) Structural model of a homodimer of ScRad50 obtained using AlphaFold2 with delimitations 1-434 and 920-1312 in blue and purple cartoons with the positions of the conserved glycine residues in contact with each other in the closed coiled-coils conformation highlighted in orange. (B) Superimposition between the *S. cerevisiae* Rad50 model shown in panel A and the cryoEM structure of the Rad50 homodimer from *Chaetomium thermophilum* (PDB: 7ZR1) highlighting the similarity and reliability of the AF2 model. (C) Predicted Alignment Error (PAE) map calculated by AlphaFold2 showing the confidence of the distances between residues of the ScRad50 model shown in panel A. (D) PAE map calculated for the model with highest AF2 confidence score for the *S. cerevisiae* Mre11 (1-533) (green) - Rad50 (1-215+1106-1312) (blue) complex generated in the first class of AF2 models in a 2:2 stoichiometry representative of an inactive form. (E) PAE map calculated for the model with highest AF2 confidence score for the *S. cerevisiae* Mre11 (1-533) (green) - Rad50 (1-215+1106-1312) (blue) complex generated in the first class of AF2 models in a 2:2 stoichiometry representative similar to an active form.

**Figure S2:**
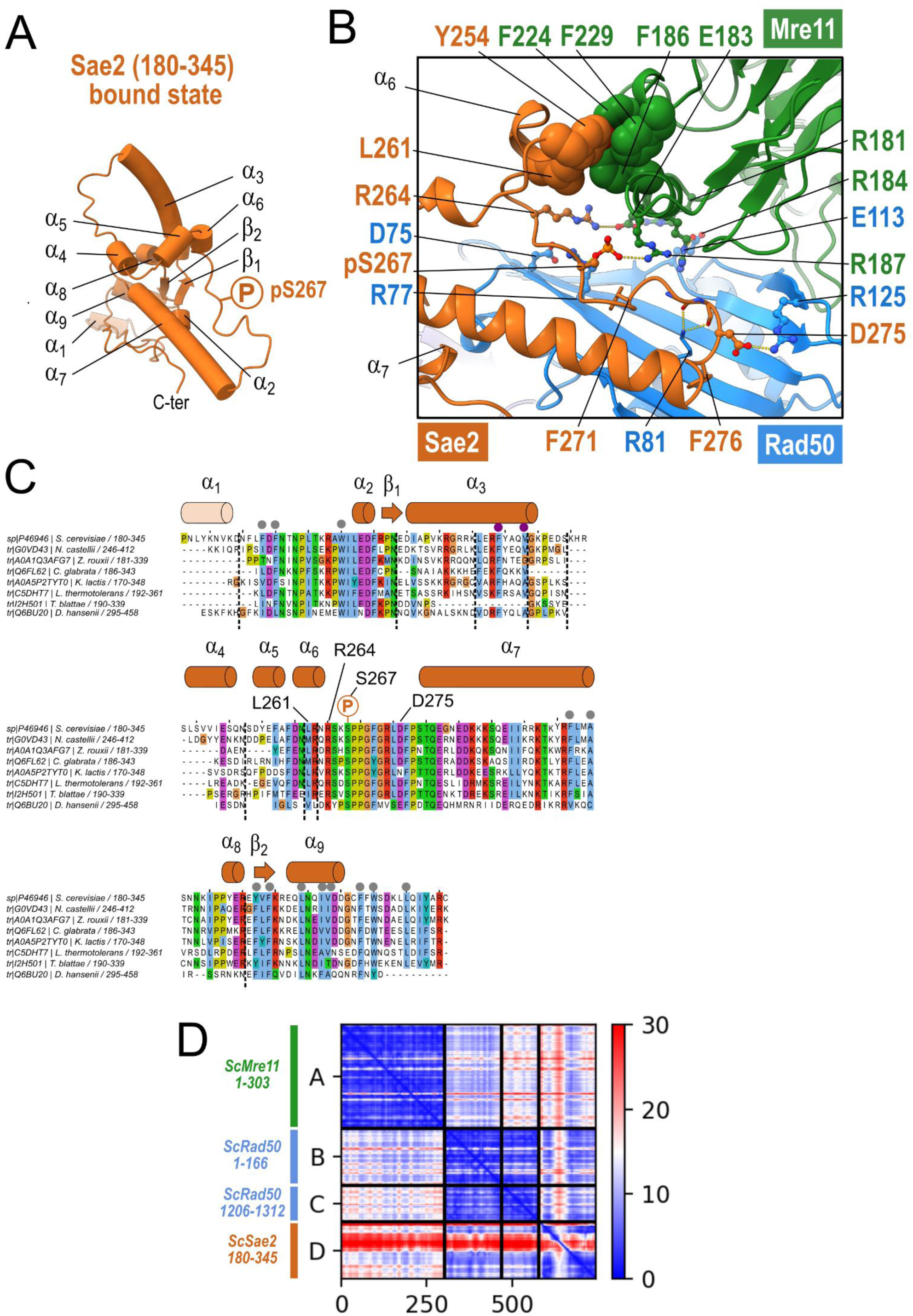
Details on the structural modeling of *S. cerevisiae* Mre11-Rad50-Sae2 assembly. Related to Figure 2. (A) Cartoon representation of the ScSae2 domain (180-345) as predicted in the bound conformation to the ScMre11 (1-303) and ScRad50 (1-166 + 1206-1312) domains. The secondary structure elements are labelled together with the position of the phosphorylated residue S267. (B) Close-up view of the structural model of the *S. cerevisiae* Sae2 (180-345) (orange) - Mre11 (1-303) (green) - Rad50 (1- 166+1206-1312) (blue) complex generated by AF2 in a 1:1:1 stoichiometry representative of the active conformation form as shown in Figure 2 with most of the amino-acids involved in the tripartite interface labelled. (C) Multiple sequence alignment of the ScSae2 domain (180-345) with closely related homologs from *Saccharomycetaceae* yeast species. Secondary structure elements as in panel A are labelled together with the residues mutated in this study. The grey dots indicate the conserved hydrophobic residues forming the hydrophobic core buried in this domain. Dotted lines represent insertion regions in the homologs of ScSae2 which are not shown for clarity (D) Predicted Alignment Error (PAE) map calculated by AlphaFold2 showing the confidence of the distances between the residues in contact in the ScMre11 (1-303) - ScRad50 (1-166 + 1206-1312) - ScSae2 (180-345) modeled complex.

**Figure S3:**
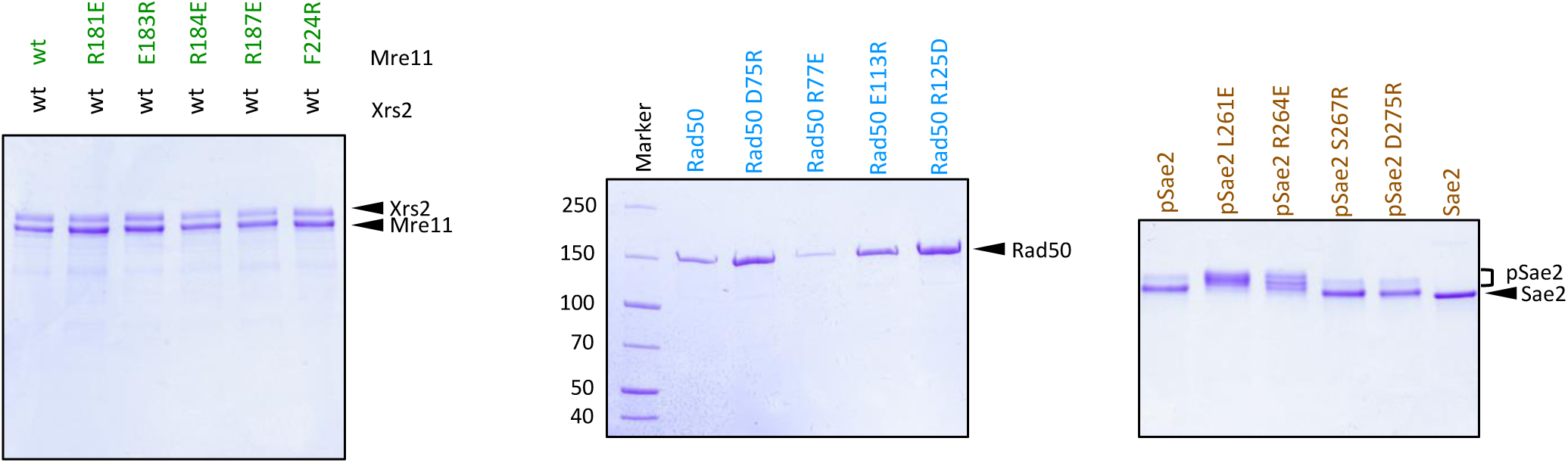
Purified recombinant proteins used in this study. Related to Figure 3. Mre11 and Xrs2 proteins were co-expressed together and purified as a complex. Rad50 variants were purified separately. Phosphorylated Sae2 and variants (indicated as pSae2) were expressed and purified in the presence of phosphatase inhibitors, resulting in an electrophoretic mobility shift. Sae2 prepared without phosphatase inhibitors (right lane, Sae2) was included as a reference. The polyacrylamide gels were stained with Coomassie blue.

**Figure S4:**
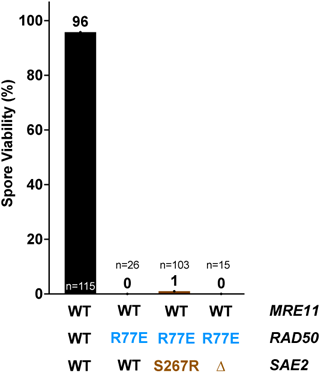
R77E substitution in Rad50 does not restore spore viability of *sae2S267R* or *sae2Δ*. Related to Figure 4. Spore viability of strains for the indicated mutations is shown. The number of tetrad dissections (n) is indicated for each.

**Figure S5:**
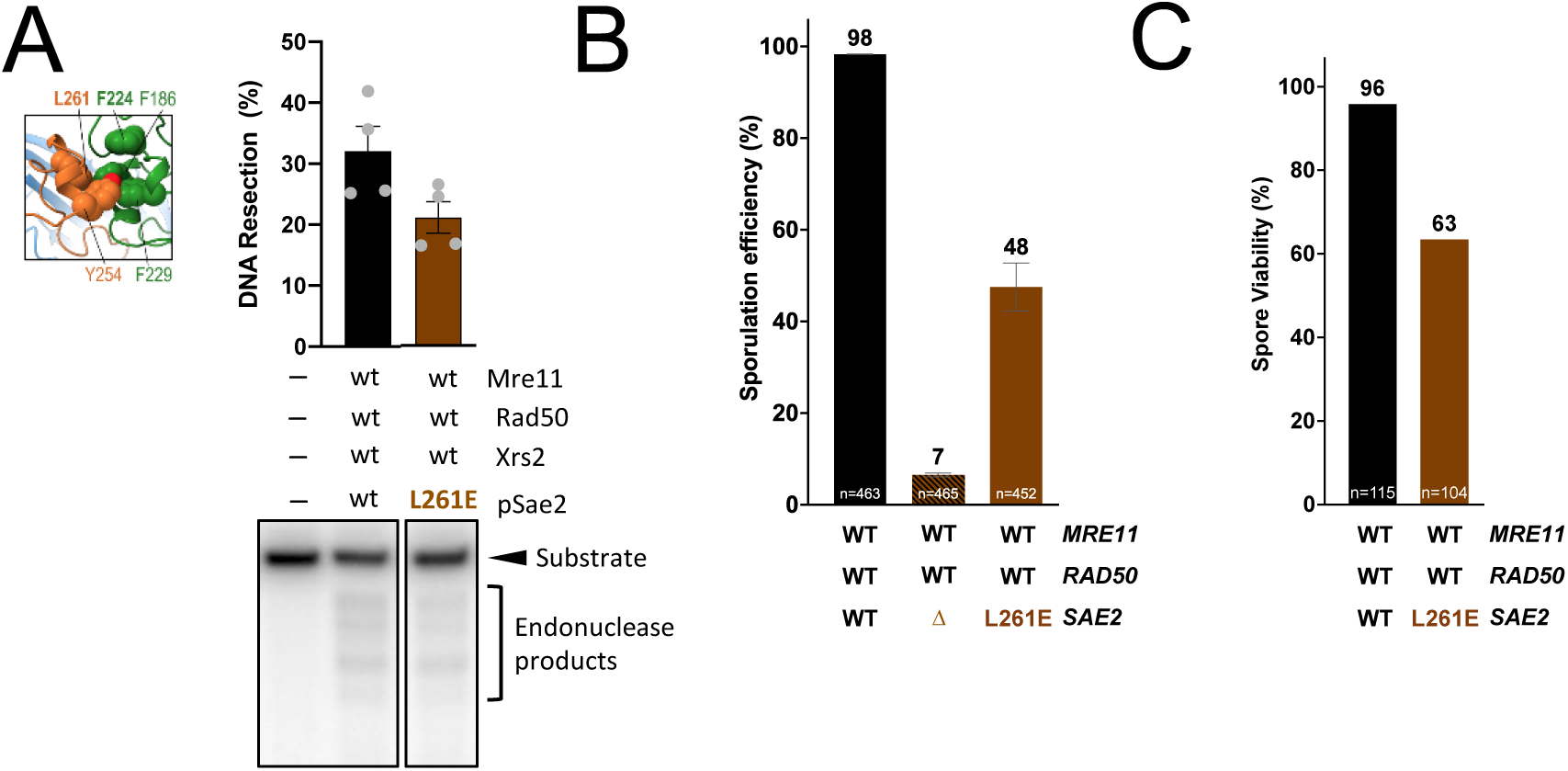
The small hydrophobic cluster between Sae2 and Mre11 favors Mre11 nuclease. Related to Figure 5. (A) Nuclease assays with the indicated variants (Mre11-Xrs2, 25 nM; Rad50, 50 nM; pSae2, 100 nM). Bottom, a representative experiment; top, a quantitation. N=3; Error bars, SEM. (B) Sporulation efficiency, as in Figure 3D. (C) Spore viability of strains with the indicated mutations. The number of tetrad dissections (n) is indicated for each.

**Figure S6:**
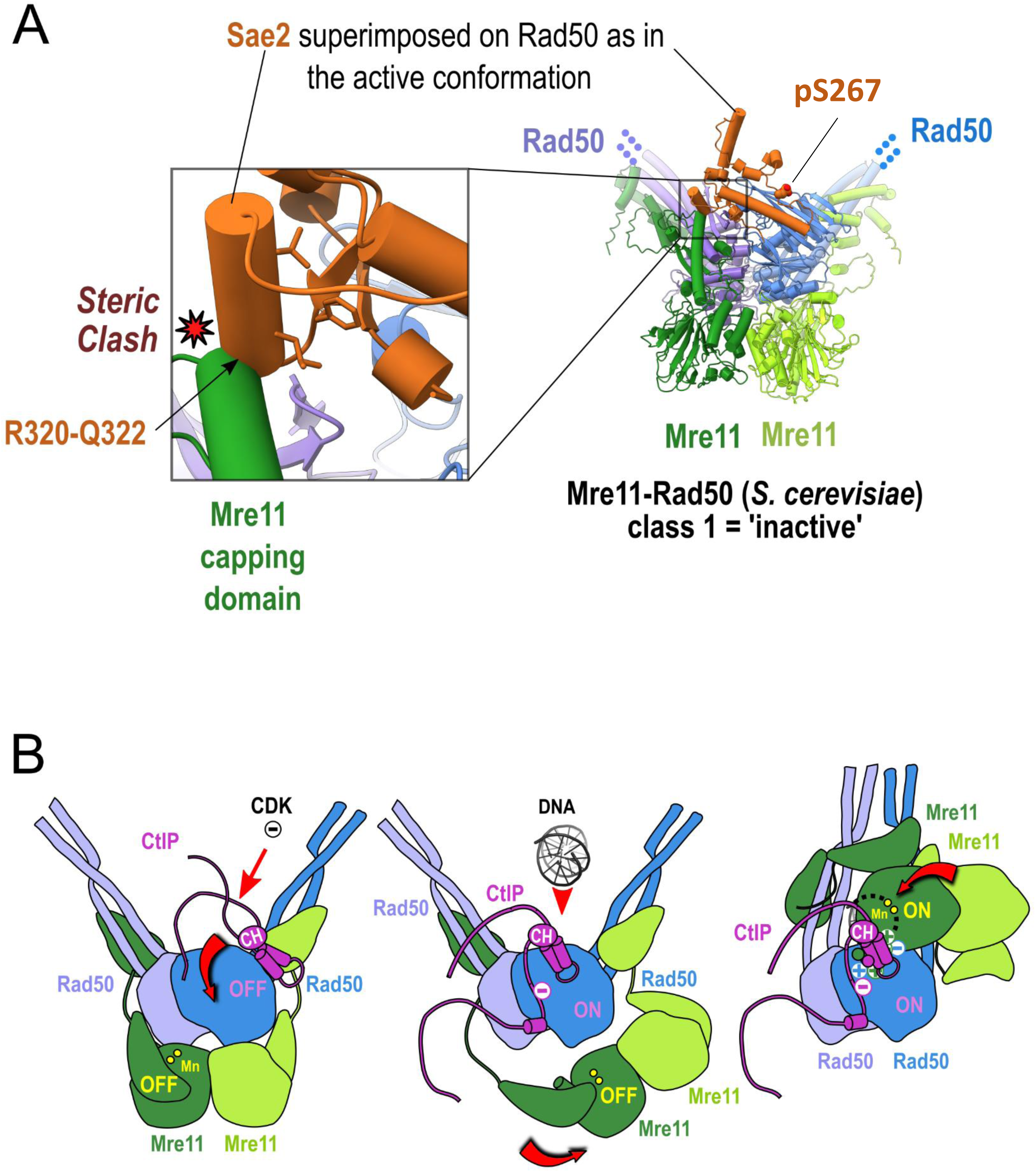
Models illustrating the molecular events occuring the transition between the inactive and active conformations. Related to Figure 6. (A) Superimposition between the structural model of the *S. cerevisiae* Mre11 (1-533) (green) - Rad50 (1-215+1106-1312) (blue) complex generated in the first class of AF2 models in a 2:2 stoichiometry representative of an inactive form and the ScSae2 domain in complex with ScMre11 and ScRad50 in the active conformation (as shown in Figure 2A) taking the outer β-sheet of Rad50 as a common reference for the superimposition. (B) Cartoon illustrating in human the transition from an inactive to an active state triggered by CtIP phosphorylation by CDK kinases on threonine 847 suggested from the structural arrangement of the CxxC/RHR motif in the predicted model from *Y. lipolytica*.

**Figure S7:**
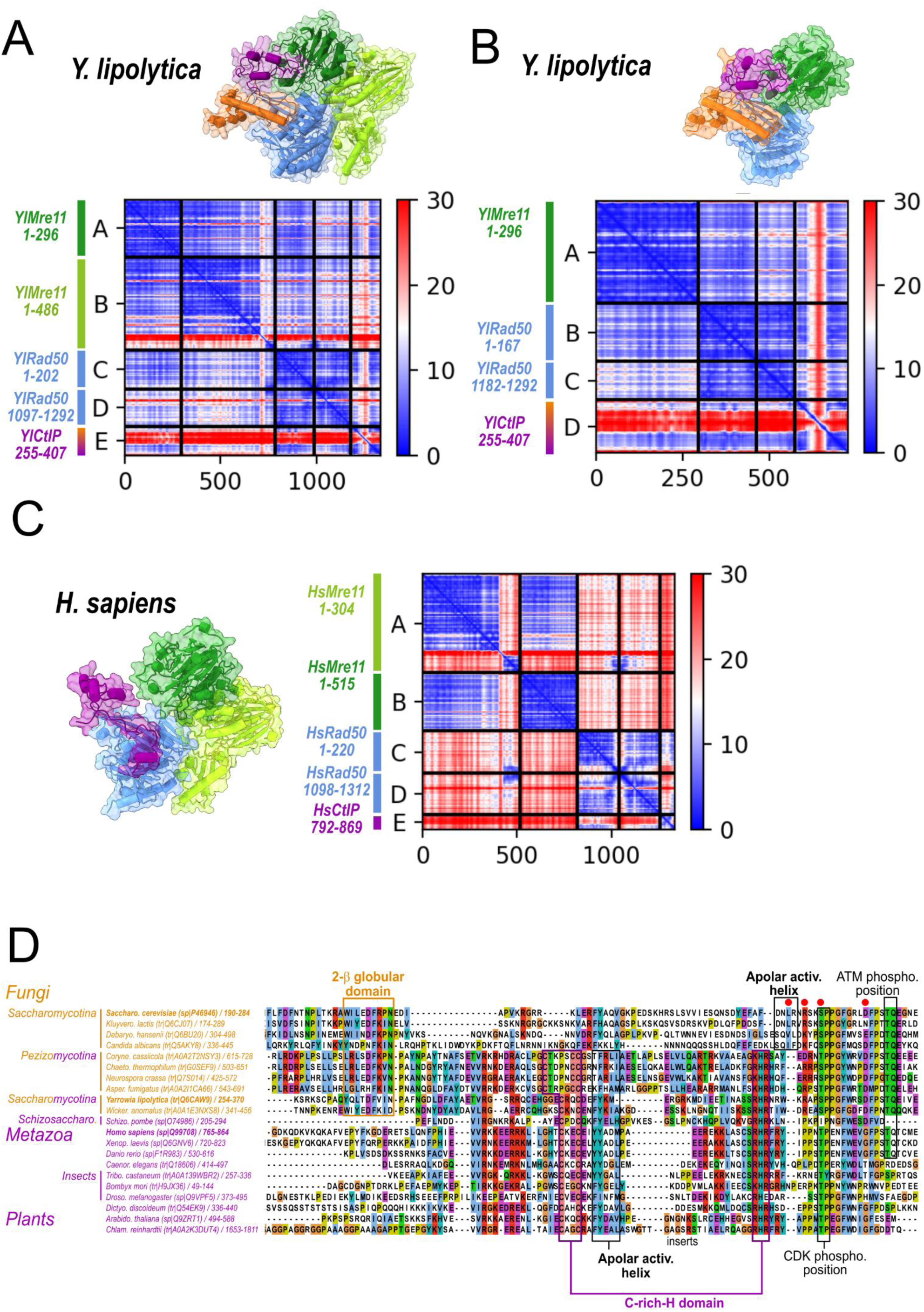
Details on the structural modeling of *Y. lipolytica* and *H. sapiens* Mre11-Rad50-Sae2 assembly. Related to Figure 7. (A) Predicted Alignment Error (PAE) map calculated by AlphaFold2 showing the confidence of the distances between the residues in contact in the *Y. lipolytica* CtIP/Sae2 (255-407) (orange+purple) - Mre11 (1-296) (green) - Mre11 (1-486) (lime) - Rad50 (1-202+1097-1292) (blue) complex generated by AF2 in a 1:1:1:1 stoichiometry representative of the active conformation form. (B) PAE map calculated by AlphaFold2 showing the confidence of the distances between the residues in contact in the *Y. lipolytica* CtIP/Sae2 (255-407) (orange+purple) - Mre11 (1-296) (green) - Rad50 (1-167+1182-1292) (blue) complex generated by AF2 in a 1:1:1 stoichiometry representative of the active conformation form. (C) PAE map calculated by AlphaFold2 showing the confidence of the distances between the residues in contact in the *H. sapiens* CtIP (792-869) (purple) – 2 Mre11 (1-304 and 1-515) (green) - Rad50 (1-220+1098-1312) (blue) complex generated by AF2 in a 1:1:1:1 stoichiometry representative of the active conformation form. (D) Multiple sequence alignment of the ScSae2 domain (190-284) a larger number of representative species than in Figure 6. Secondary structure elements as in panel A are labelled together with the residues mutated in this study. The grey dots indicate the conserved hydrophobic residues forming the hydrophobic core buried in this domain. Dotted lines represent insertion regions in the homologs of ScSae2 which are not shown for clarity

### Supplementary Tables legends

**Table S1:**
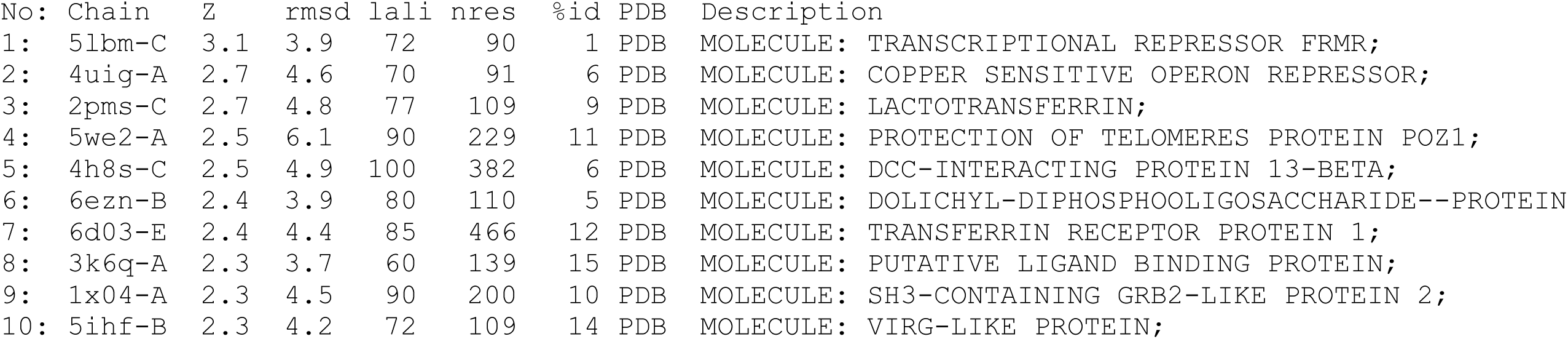
Ten top hits from the DALI server analysis.

**Table S2:**
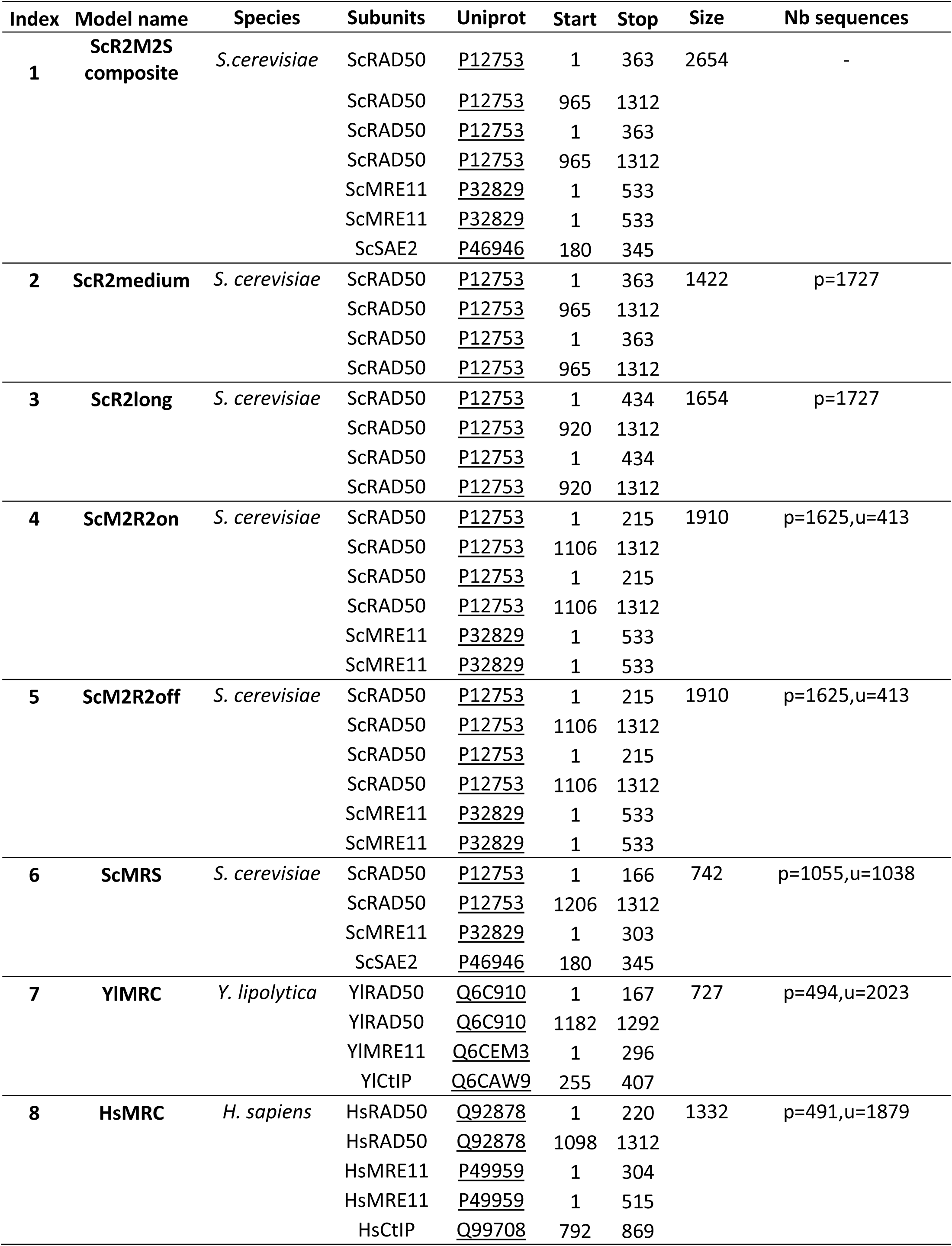
Description of conditions used to generate models.

**Table S3:** Summary of evaluation scores for models generated by AlphaFold2.

| Index | Model name | Species | pLDDT | pTMscore | ipTMscore | confidence_score | PAE |
| --- | --- | --- | --- | --- | --- | --- | --- |
| 2 | ScR2medium | *S. cerevisiae* | 79.8 | 0.54 | 0.51 | 0.52 |  |
| 3 | ScR2long | *S. cerevisiae* | 81.2 | 0.48 | 0.46 | 0.46 | Fig. S1C |
| 4 | ScM2R2on | *S. cerevisiae* | 81.2 | 0.68 | 0.65 | 0.66 | Fig. S1E |
| 5 | ScM2R2off | *S. cerevisiae* | 85.5 | 0.87 | 0.86 | 0.86 | Fig. S1D |
| 6 | ScMRS | *S. cerevisiae* | 84.7 | 0.75 | 0.69 | 0.70 | Fig. S2D |
| 7 | YIMRC | *Y. lipolytica* | 83.5 | 0.8 | 0.76 | 0.77 | Fig. S7B |
| 8 | HsMRC | *H. sapiens* | 83.7 | 0.71 | 0.63 | 0.65 | Fig. S7C |

**Table S4:** Links to the ModelArchive database containing model coordinates, multiple sequence alignments and evaluation scores.

| Index | Model name | ModelArchive |
| --- | --- | --- |
| 1 | ScR2M2S composite | https://modelarchive.org/doi/10.5452/ma-jf8hv |
| 2 | ScR2medium | https://www.modelarchive.org/doi/10.5452/ma-i9sv6 |
| 3 | ScR2long | https://modelarchive.org/doi/10.5452/ma-vje4m |
| 4 | ScM2R2on | https://modelarchive.org/doi/10.5452/ma-jfoh4 |
| 5 | ScM2R2off | https://modelarchive.org/doi/10.5452/ma-lbblz |
| 6 | ScMRS | https://www.modelarchive.org/doi/10.5452/ma-qhmpy |
| 7 | YIMRC | https://www.modelarchive.org/doi/10.5452/ma-ks31s |
| 8 | HsMRC | https://www.modelarchive.org/doi/10.5452/ma-qtbol |

**Table S5:**
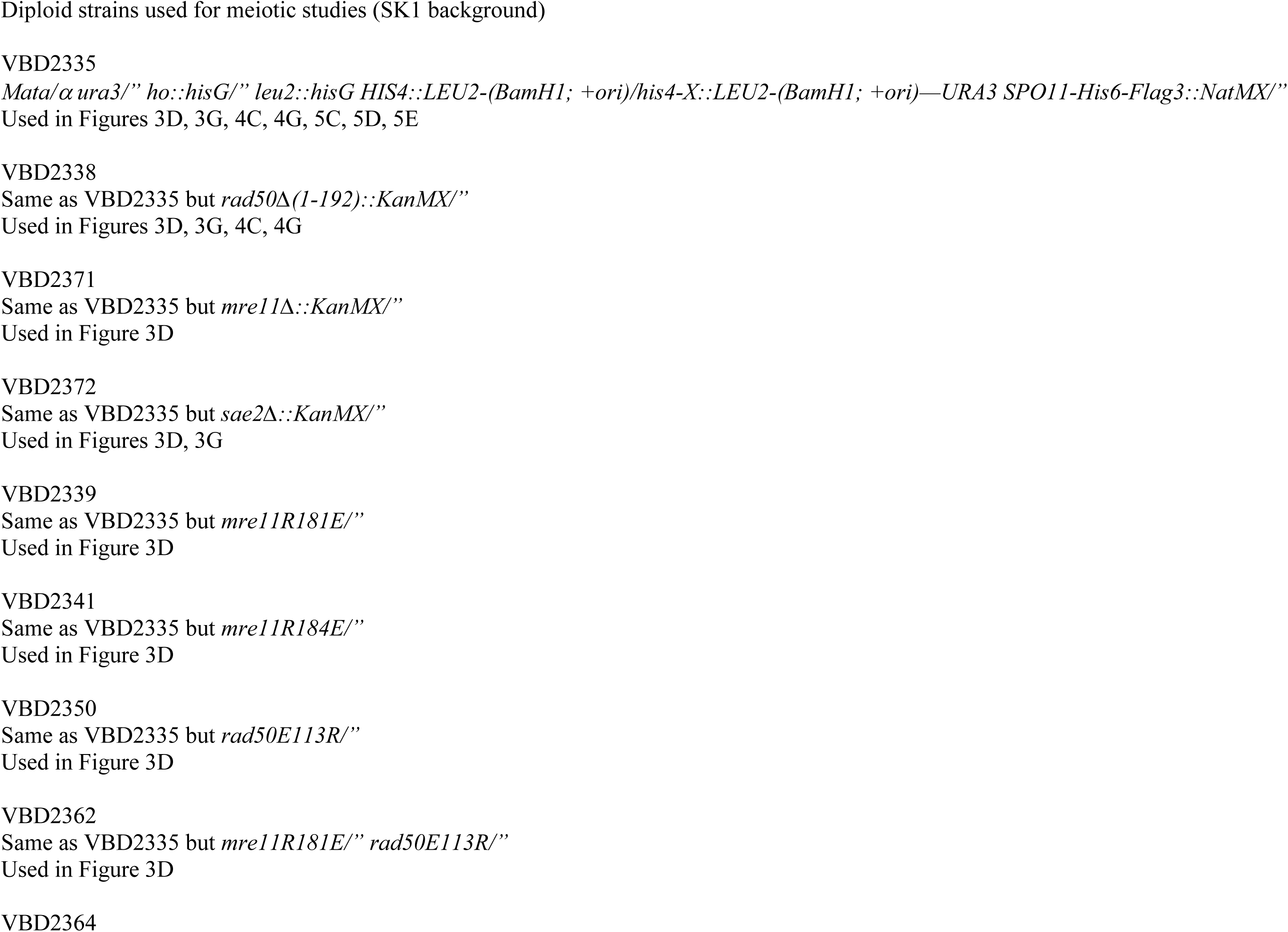
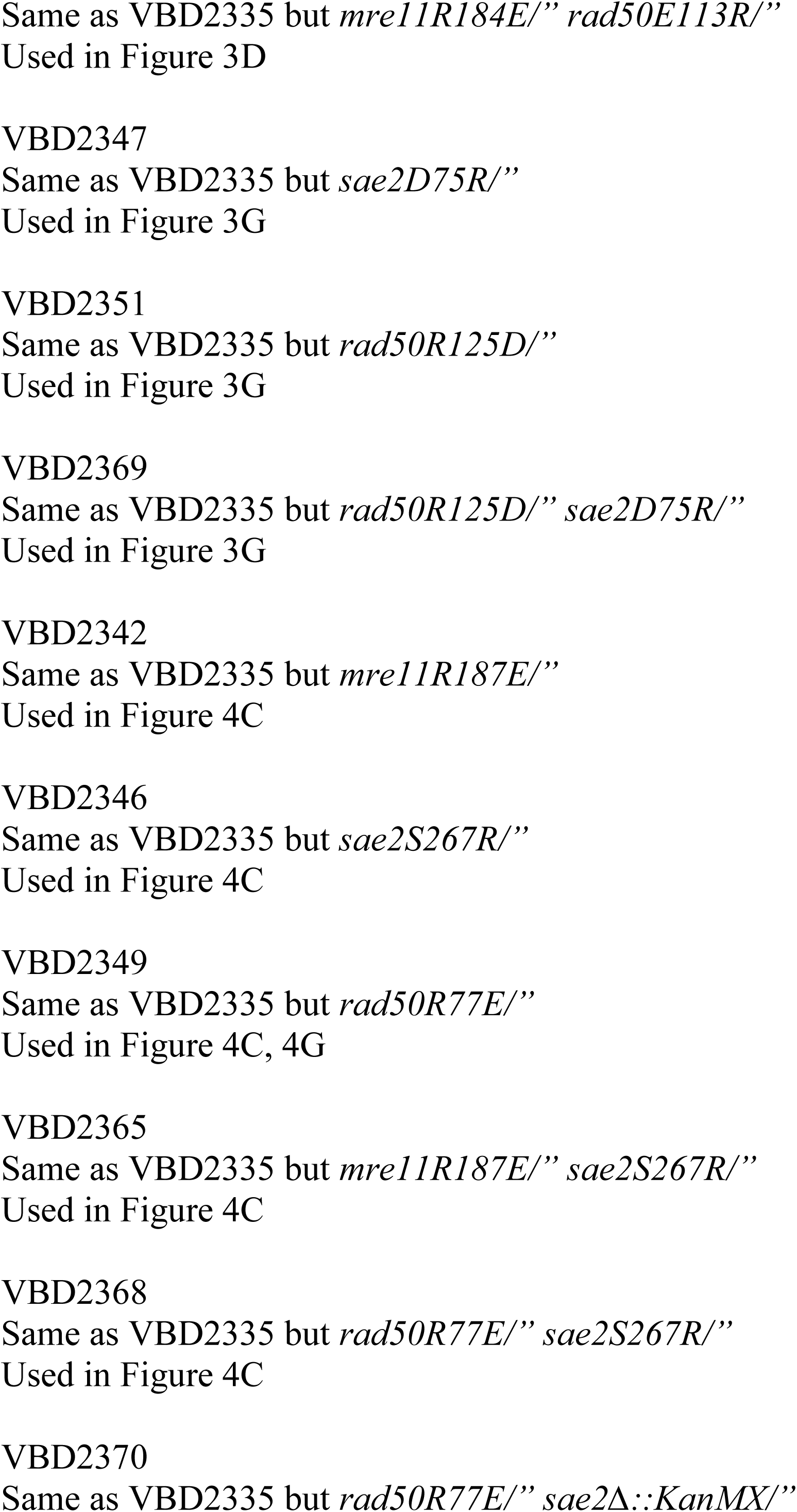
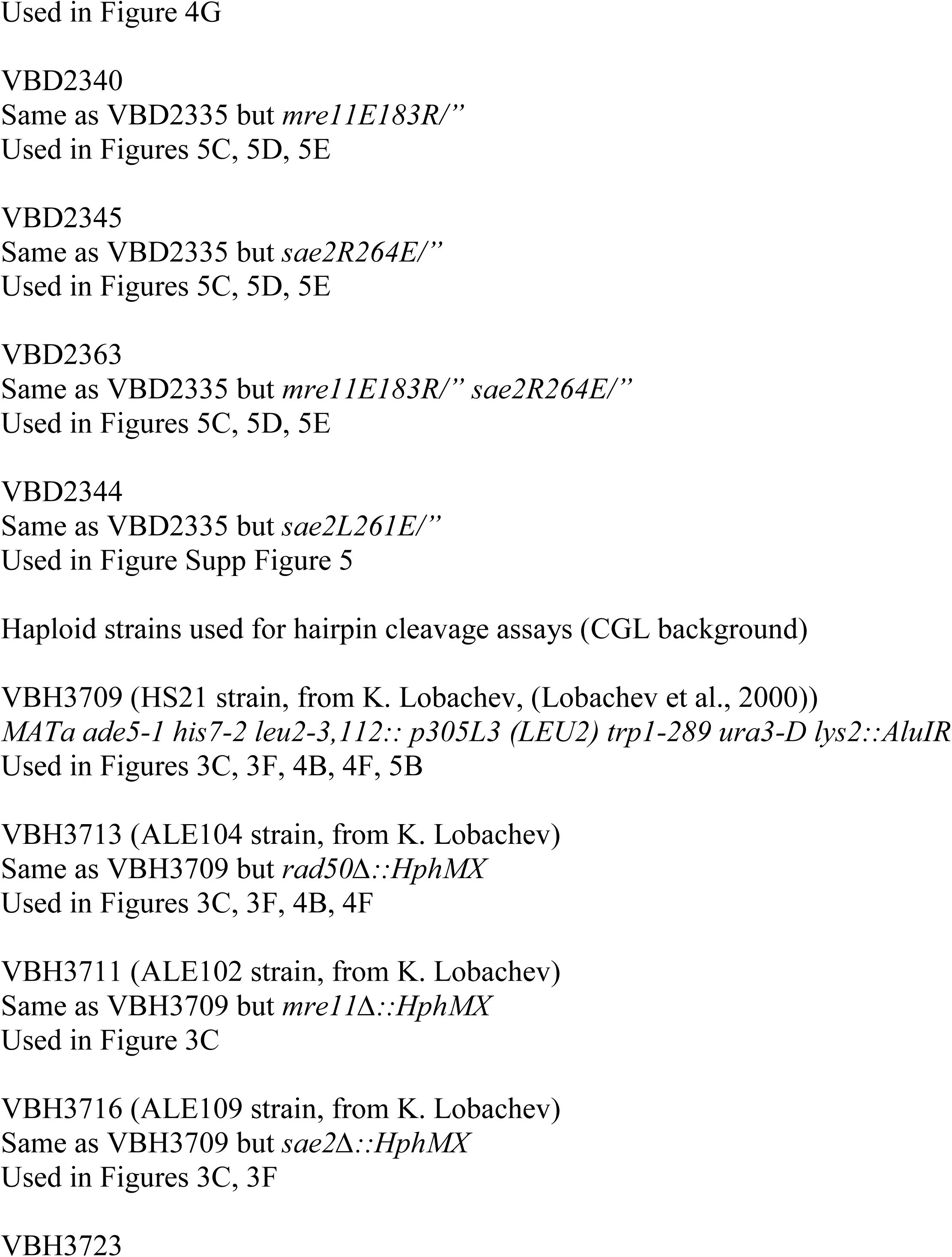
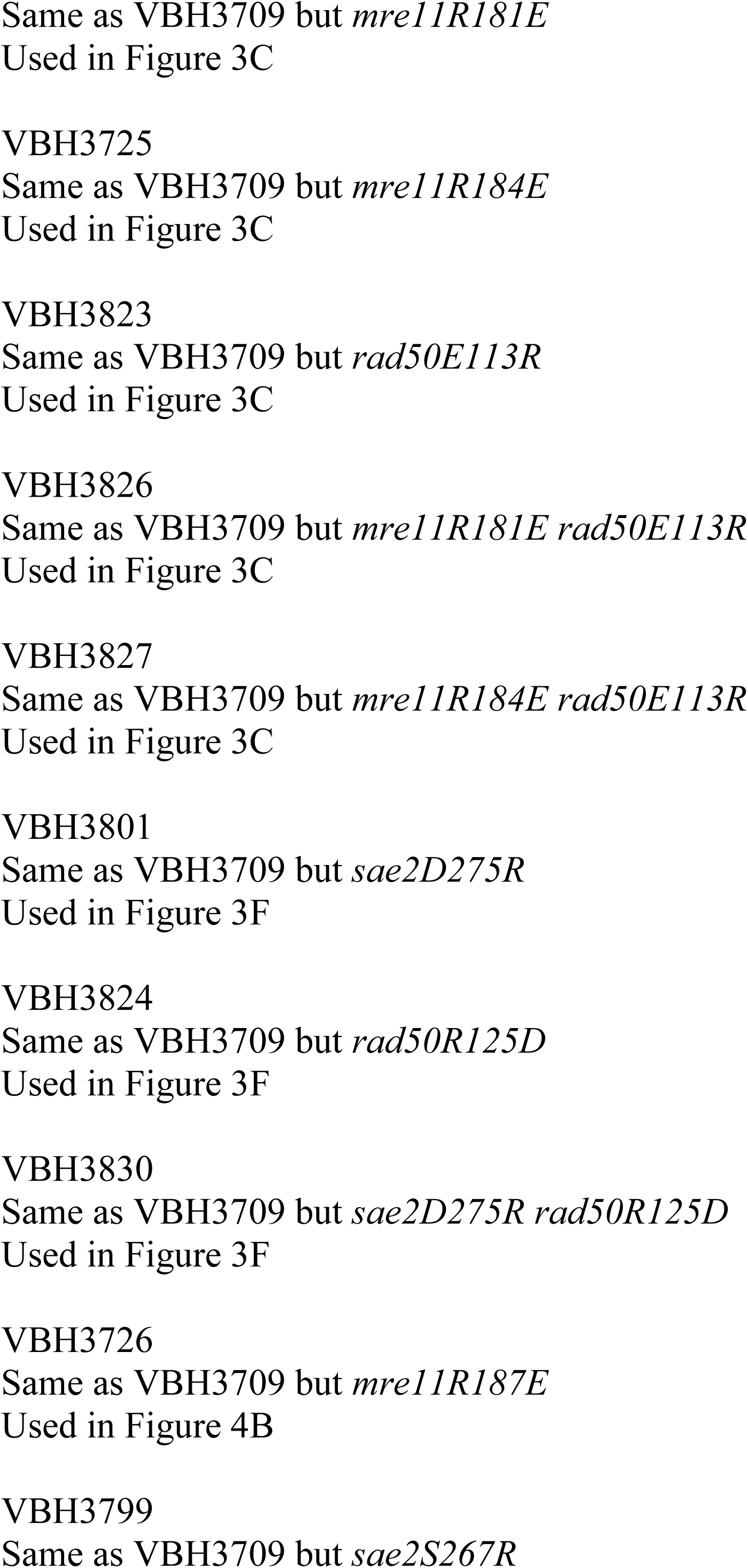
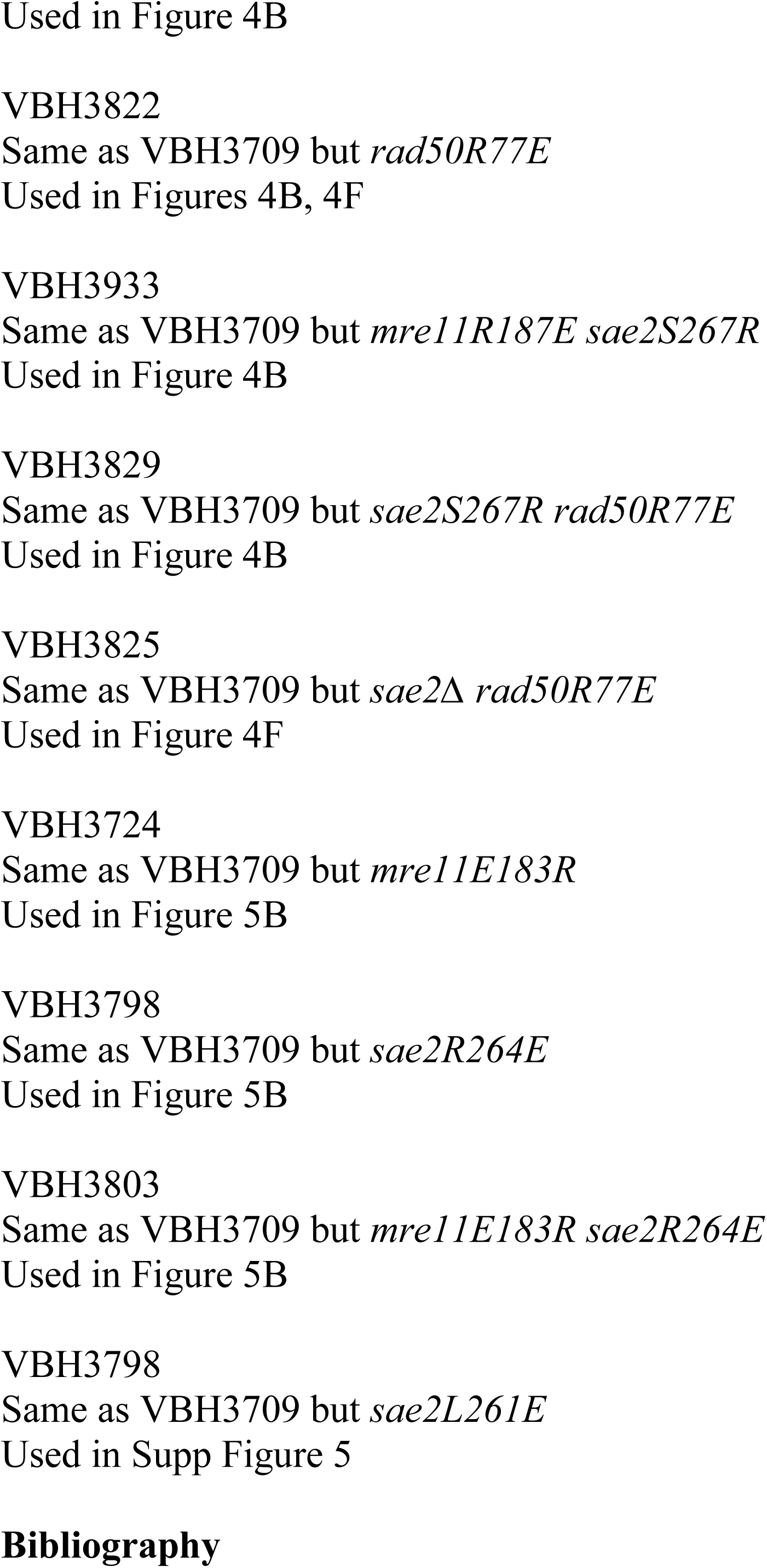
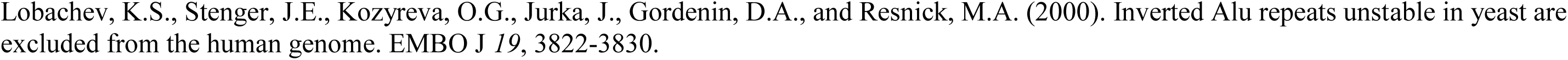
Yeast strains used in this study.

